# Computational design of potent, broadly neutralizing anti-Nipah virus and Hendra virus miniproteins

**DOI:** 10.64898/2026.09.14.751546

**Authors:** Jeremiah N. Sims, Risako Gen, Kaitlin Sprouse, Young-Jun Park, Zhaoqian Wang, Moushimi Amaya, Alka Jays, Brendan B. Larsen, Cameron Stewart, Jack T. Brown, Robert Ragotte, Rashmi Ravichandran, Garrett Ruth, Xinting Li, Dionne Vafeados, Jesse D. Bloom, Christopher Broder, David Veesler, David Baker

## Abstract

The prototype members of the genus *Henipavirus,* Nipah virus (NiV) and Hendra virus (HeV), cause recurrent zoonotic spillovers with case fatality rates ranging from 40-90% in humans. Currently, there are no approved vaccines or therapeutics for use in humans. Neutralizing antibodies targeting the NiV/HeV F-or G-glycoproteins protect animals from lethal challenge and are a main correlate of protection. However, antibody-based formulations are expensive, typically requiring hospital admission for administration and cold-chain for storage and transportation. To address the lack of shelf-stable clinical countermeasures, we computationally designed thermostable miniproteins that cross-react with subnanomolar affinities with both NiV and HeV F and G glycoproteins and inhibit viral entry *in vitro* with potencies comparable to lead antibodies. Oligomerized forms of these miniproteins have enhanced potency relative to their monomeric building blocks and increase the barrier for emergence of escape mutants, establishing them as promising preclinical candidates against these deadly viruses.

## Main

Nipah virus (NiV) and Hendra virus (HeV) are zoonotic bat-borne paramyxoviruses, in the genus *Henipavirus* (HNV), that are responsible for recurrent outbreaks of lethal disease in South & Southeast Asia, and Australia. Using pteropid bats as a reservoir, these viruses can infect a variety of mammalian hosts, most notably livestock (horses, pigs, etc.) that facilitate their transmission to humans via fluid contact or respiratory routes (Luby and Gurley 2012; Weatherman et al. 2018). NiV and HeV are the only recognized henipaviruses that can cause “henipaviral disease”, a severe respiratory and/or neurological pathogenesis, with case fatality rates ranging from 40-90% in humans (Kenmoe et al. 2019; Vasudevan et al. 2024; Satter et al. 2023). HeV appears restricted to Australia, and although known NiV outbreaks have occurred in Malaysia, Singapore, India, Bangladesh, and the Philippines, serological evidence of NiV or Nipah-like viruses in bats is evident in multiple additional countries, including, Cambodia, China, Indonesia, Madagascar, Thailand, and VietNam, indicating that a large fraction of the global population is at risk of spillovers (Moore et al. 2024). There are no approved vaccines or specific therapeutics for use in humans, though clinical trials are currently ongoing or planned for two monoclonal antibodies, m102.4 & hu1F5, inhibiting viral entry by targeting the surface glycoproteins (Playford et al. 2020; Zeitlin et al. 2024).

NiV and HeV entry into cells is mediated by two surface glycoproteins working in concert to promote attachment to host receptors (G) and fusion of the viral and host membranes (F). The HNV G protein is a tetrameric type-II integral membrane protein that features a receptor-binding, beta-propeller head domain that binds with high affinity to its cognate human receptor, Ephrin-B2 (EFNB2), as well as EphrinB3 albeit with lower affinity (Bonaparte et al. 2005; Narayanan et al. 2023b; Z. Wang, Amaya, et al. 2022; Larsen et al. 2025; Negrete et al. 2005). HNV G receptor engagement is believed to trigger conformational changes propagated to the trimeric HNV F protein to initiate membrane fusion (Q. Liu et al. 2013). Antibodies inhibiting attachment and fusion correlate with protection in animal models and have been a primary focus of therapeutic development (Mire et al. 2020; Z. Wang, Dang, et al. 2022; Doyle et al. 2021; Zhu et al. 2008; Xu et al. 2013; Geisbert et al. 2014; Playford et al. 2020; Zeitlin et al. 2024; Dang et al. 2019, 2021).

Antibody m102.4 inhibits receptor binding to NiV and HeV G competitively, and protects Syrian hamsters, ferrets, and African green monkeys from lethal disease (Dong et al. 2020; Zeitlin et al. 2024; Bossart et al. 2009). However, m102.4 proved less effective than anti-NiV/HeV F antibody hu1F5 in African green monkeys against the NiV Bangladesh strain (NiV-B), a highly virulent strain, despite comparable efficacies *in vitro*. This is thought to stem from valency mismatch – the NiV G tetramer maintains open binding sites when a bivalent antibody binds, an effect that may be mitigated by engineering a tetravalent competitive inhibitor (Zeitlin et al. 2024). In the same study, hu1F5, a monoclonal antibody that inhibits fusion through binding at a highly conserved quaternary epitope of the HNV F protein, protected animals against all tested strains of HNV *in vivo*. These antibody-based therapeutics have potential to be potent additions to the pandemic preparedness toolbox, owing to their distinct mechanisms of neutralization and m102.4 has already been administered to multiple individuals on a compassionate basis. However, antibody-based therapeutics are expensive to manufacture and deploy at scale, which reduces their utility in resource-limited settings (Sifniotis et al. 2019).

We reasoned that computational protein design could generate anti-NiV/HeV G and F proteins with improved stability and potency relative to mAbs, and that the modularity of design could enable exploration of oligomerization states matching that of the viral targets. Here, we describe the design of highly stable HNV F-and G-targeted proteins, and their multimerization to form oligomers maximizing neutralization potency and breadth.

### Design of G-directed minibinders interfering with receptor engagement

We first set out to design binders that prevent HNV G from binding to its cellular receptor hEphrinB2 (EFNB2) by embedding the interacting elements from EFNB2 into a *de novo* designed scaffold. Starting from a crystal structure of the complex of EFNB2 and NiV G (PDB 2VSM), the EFNB2 G-H loop and β-strand (residues 109-128, QDIKFTIKFQEFSPNLWGLE) were chosen as a minimal motif to scaffold (**Figure 1a-b**). Previous research supports the importance of the G-H loop for binding to HNV G (Narayanan et al. 2023a; Bowden et al. 2008), and we hypothesized that the adjacent β-strand itself would serve as an essential secondary structural feature to guide RFdiffusion to create a soluble scaffold to support this motif. A total of 18,933 backbones were generated, which then served as input for sequence design and folding. Initially, the designs were folded with AlphaFold2 (AF2) in the context of templated NiV G (i.e. NiV G structure was given to the model) to evaluate confidence of folding (pLDDT), likelihood of complexing with NiV G (pAE_interaction – interchain predicted aligned error), and structural agreement with the diffused backbone (C_α_-RMSD). However, AF2 failed to recapitulate the G-H loop in the pocket, resulting in pAE_interaction scores > 25Å and folding predictions placed distantly from the NiV G target (**Figure 1c – purple**). A range of pAE_interaction values (5-28Å) was obtained when the loop interface was forced by templating consecutive G-H loop pocket residues “F-S-P-N” (**Figure 1c – teal**), permitting *in silico* discrimination between predicted binders and non-binders using a standard pAE_interaction cutoff of <10 Å (Watson et al. 2023). After filtering the loop-templated predicted structures based on prediction quality (pLDDT > 80, pAE_interaction < 10 Å, C_α_-RMSD < 2Å), 24/37,873 designs for which AF2 structure predictions matched the design model were selected for experimental characterization (see Methods).

**Figure 1.**
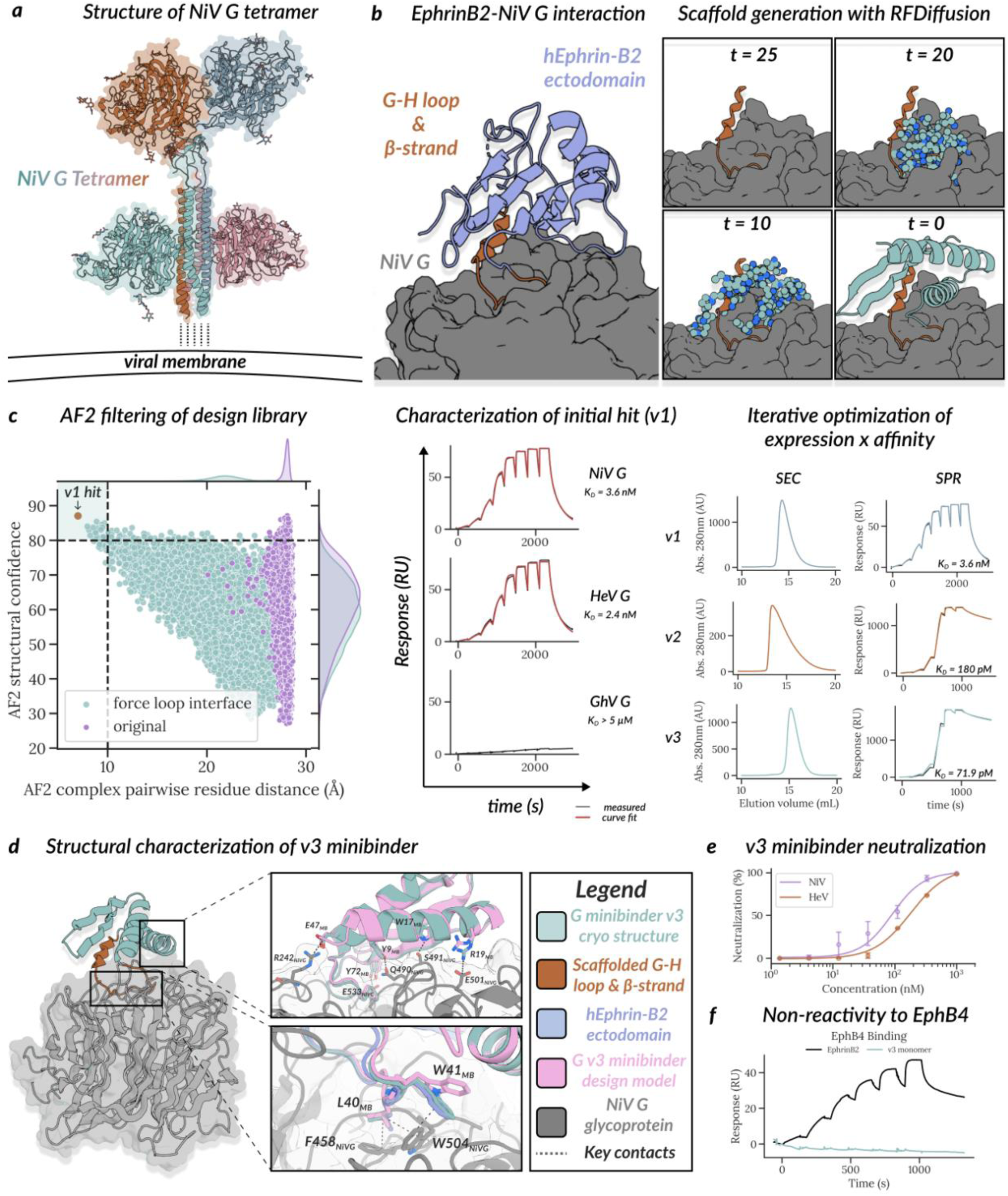
Computational design of picomolar HNV G minibinders blocking receptor engagement. **(a)** Cryo-electron microscopy (Cryo-EM) structure of the Nipah virus (NiV) G protein ectodomain tetramer (composite of PDBs: 7TY0 and 7TXZ). The bound nAH1.3 Fab fragments are not shown for clarity. **(b)** (Left) Crystal structure (PDB: 2VSM) of the hEphrinB2 (hEFNB2, purple) ectodomain bound to the NiV G head (grey). The EFNB2 G-H loop (orange) was chosen for scaffolding because it constitutes the majority of the EFNB2-NiV G interface and is essential for HNV G attachment. (Right) Snapshots of the RFDiffusion trajectory that yielded G minibinder v1 (teal), which harbors the EFNB2 G-H loop (orange). **(c)** (Left) Selection of anti-HNV G designs based on AF2 (pLDDT) and AF2 pairwise alignment error (pAE_interaction). We could only achieve accurate binding predictions (RMSD < 2Å) with a meaningful distribution of pAE_interaction values (AF2 metric for binding propensity, 5-28Å) by templating 4 residues (FSPN) of the G-H loop. This allowed for downselection of the library to 24 wet lab candidates by high structural confidence and predicted binding mode to NiV G and HeV G. *In silico* hits were recombinantly expressed, purified, and then tested via competition BLI using immobilized NiV G as ligand and EphrinB2 as analyte in the presence of 10µM minibinder (see Supplementary Figure 2). (Middle) Surface plasmon resonance (SPR) single-cycle kinetics evaluation of binding of G minibinder v1 (titrated 5x over 6 steps from 5 µM to 1.6 nM) to immobilized NiV G and HeV G head domains. Curve fit (red) to measured signal (black) was used to determine kinetic and equilibrium binding constants. The v1 hit was then optimized for both expression yield and affinity to achieve a 50x improvement in affinity (3.6nM → 71.9 pM) **(d)** Cryo-EM structure of G minibinder v3 (teal) bound to the NiV G head (grey) determined to 2.7 Å resolution. The HENV-32 Fab used to assist structural determination was omitted for clarity. The Ephrin-B2 G-H loop adopts a similar conformation to the v3 minibinder’s scaffolded G-H loop in the NiV G pocket. **(e)** *In vitro* neutralization of NiV F/G-and HeV F/G-harboring recombinant CedV chimeras by G minibinder v3. **(f)** Evaluation of binding specificity of the G minibinder v3 which does not interact with hEphB4, the native EFNB2 binding partner.

Gene fragments encoding the 24 designs (all ∼10 kDa) were expressed with a C-terminal tandem SNAC tag and 6x-His tag and purified with NiNTA and size exclusion chromatography (SEC) as previously described (Watson et al. 2023). Out of the screened designs, 18/24 expressed, and 6/24 had prominent peaks in the expected monomer fraction (2/22 monodisperse) between 14-16 mL (**Supplementary Figure 1**). Using biolayer interferometry, we first evaluated competitive binding by immobilizing EFNB2 on BLI biosensors and assessing binding to tetrameric NiV G in the presence and absence of 100x molar excess of the G minibinder v1 designs and identified G minibinder v1.10 as a hit that reduced NiV G binding to Ephrin-B2. We followed up with immobilizing the hit to BLI biosensors and titrated against NiV G as analyte (2x dilutions from 100nM to 1.56nM) and observed dose-dependent increases in on-rate with increasing analyte concentration (**Supplementary Figure 2**). Using surface plasmon resonance (SPR) to interrogate kinetic and equilibrium binding constants, we found that G minibinder v1 not only bound tightly to immobilized NiV G (K_D_=3.6 nM) but also cross-reacted with HeV G (K_D_=2.4 nM). No binding was detected to Ghanaian bat virus G, a distantly-related henipavirus sharing only 26-29% sequence similarity to NiV G and HeV G (**Figure 1c**).

Given that the tight binding of the cognate EFNB2-NiV G complex (∼340 pM) requires a high affinity inhibitor for competition-based neutralization, we aimed to improve the affinity of the G minibinder v1 using partial diffusion and ProteinMPNN (Watson et al. 2023) by iteratively peturbing the backbone and redesigning its sequence. In the first round, we tested 64 designs at 4-mL culture scale (Watson et al. 2023), and identified a G minibinder v2 that had a prominent peak at the expected elution volume by SEC and improved affinity (K_D_ = 0.176 nM, **Figure 1c**). Given the propensity of G minibinder v2 to aggregate, we used SolubleMPNN to redesign its core and solvent-exposed residues (outside the interface) to yield G minibinder v3. Of the 35 updated designs tested, 8 were monomeric and monodisperse by SEC and 3 bound NiV G with approximate affinities < 200 pM. G minibinders v3 and v3.23, were selected for further characterization as they were highly soluble, expressed at high levels, and had high affinity to both NiV G (69 pM & 105 pM, resp.) and HeV G (72 pM & 81 pM, resp., **Figure 1c**). Despite sharing 86% sequence identity, G minibinder v3 was selected owing to higher affinities relative to v3.23-based constructs (**Supplementary Figure 3**).

We determined a cryo-EM structure of the NiV G head domain in complex with G minibinder v3 (and the HENV-32 Fab to aid structural determination) at 2.7 Å resolution, which shows that the cognate EFNB2 loop-strand interface is recapitulated upon binding (**Figure 1d, Supplementary Figure 4**). The structure closely resembles the designed model (minibinder C_α_-RMSD = 1.75 Å when superimposing the NiV G head domain), including the scaffolded region of EFNB2 from PDB 2VSM (C_α_-RMSD = 1.36 Å) and the G-H motif interactions with the NiV G pocket. An average surface area of 1,420 Å is buried at the interface between the minibinder and the G head domain through shape complementarity and polar interactions. L40_MB_ forms extensive van der Waals interactions with F458_NiVG_ and W504_NiVG_,W41_MB_ forms T-shaped pi-stacking interactions with W504_NiVG_, whereas the neighbouring residues E47_MB_ and Y72_MB_ interact with R242_NiVG_ and E533_NiVG_ via salt bridge and hydrogen bonding, respectively. Moreover, Y9_MB_, R19_MB_, and W17_MB_ respectively form hydrogen bonds with Q490_NivG_, E501_NivG_, and S491_NiVG_. The G minibinder v3, therefore, recapitulates many interactions formed by EFNB2 with NiV G and also makes additional interactions not seen with the cognate receptor.

We next tested the ability of the G minibinder v3 to inhibit viral entry using replication-competent recombinant Cedar virus (rCedV) expressing GFP and harboring the NiV-B or HeV F and G glycoproteins instead of the endogeneous CedV counterparts. The minibinder was titrated and co-incubated with either rCedV-NiV-B-GFP or rCedV-HeV-GFP prior to addition to Vero 76 cells and fluorescent foci were counted after 24h. The G minibinder v3 was shown to block viral entry with an IC_50_ of 90.54 nM for NiV-B and 182.48 nM for HeV (**Figure 1e**). Additionally, we tested whether G minibinder v3 would bind EFNB2’s cognate partner EphB4, a receptor tyrosine kinase that plays a role in vascular development. To this end, we used SPR to measure the affinity of G minibinder v3 and soluble recombinant hEFNB2 to immobilized recombinant EphB4 (ACROBiosystems). While robust binding to EphB4 was observed for EFNB2, no binding was detected with G minibinder v3 (**Figure 1f**), indicating that this miniprotein is unlikely to stimulate off-target signaling through this receptor despite sharing an interface with EFNB2.

Collectively, these data show that G minibinder v3 is a NiV/HeV G cross-reactive picomolar binder that competitively blocks receptor engagement without interacting with hEphB4.

### Design of F-directed minibinders interfering with membrane fusion

Prior work showed that a bivalent, biparatopic anti-HNV antibody was endowed with superior neutralization potency than the parent monoparatopic, bivalent antibodies while also conferring resistance to emergence of viral escape mutants (Isaacs, Nieto, Zhang, Modhiran, Barr, Thakur, Low, Parry, Barnes, Jara, Himelreichs, Yao, Deride, Barthou-Gatica, Salinas-Rebolledo, Pamela, et al. 2025). We therefore sought to design miniproteins blocking F protein function, reasoning that these could be coupled to anti-G inhibitors to enhance neutralization potency and minimize potential for neutralization escape through viral evolution. We set out to design binders to the NiV antigenic site recognized by two neutralizing antibodies, hu1F5 and hu5B3, that bind overlapping epitopes that are highly conserved between NiV F and HeV F (**Figure 2a**). We used the 5B3-bound F cryo-EM structure (PDB 6TYS) to select target hotspots to guide the generation of 54,873 initial backbones using RFDiffusion. Of note, these miniprotein backbones spanned a tertiary epitope over just one protomer, a subset of the larger quaternary epitope of 5B3 conferred by its heavy and light chains. Sequences for each backbone were subsequently designed with ProteinMPNN before predicting the resulting complex structures with AlphaFold2 for evaluation. The best performing backbones were subjected to partial diffusion with RFDiffusion and subsequent sequence design and folding; 1601 designs with miniprotein pLDDT ≥ 85, pAE_interaction < 10, and Rosetta ΔΔG score < -40, ranging from 55 to 85 residues, were selected for experimental characterization (**Figure 2b**).

**Figure 2.**
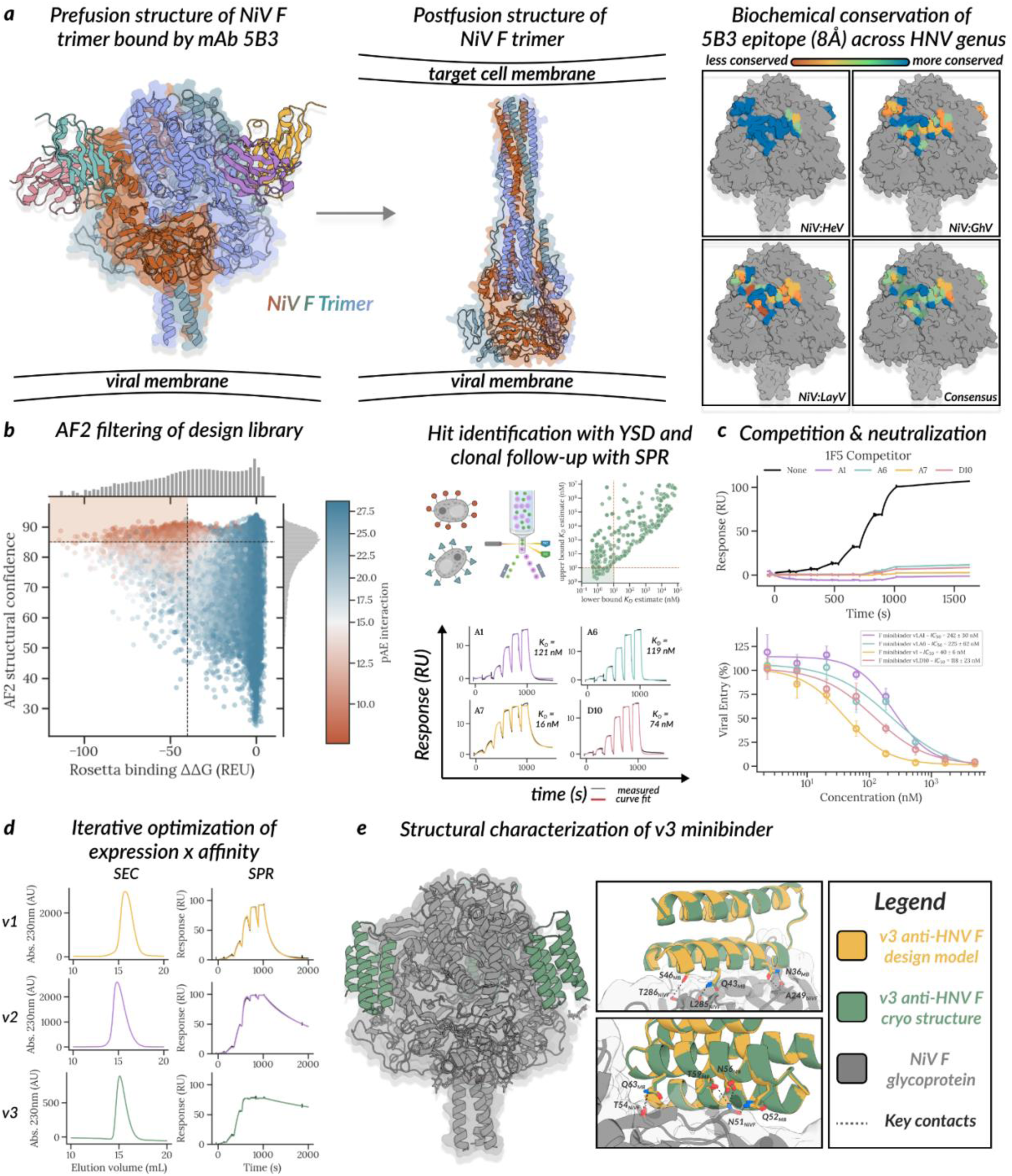
Design and characterization of anti-HNV F minibinders. **(a)** Ribbon diagram of NiV F in the mAb 5B3-bound prefusion state **(left, PDB: 6TYS)** and post-fusion **(middle, AF2 prediction)** conformation positioned relative to the viral membrane. A subset of the 5B3 quaternary epitope was targeted due to its high sequence conservation among NiV and HeV, portrayed as all residues within 8 Å of the Fab **(right)**. Sequence conservation at this epitope decreases for more distant relatives GhV and LayV. **(b)** ProteinMPNN and RFDiffusion designs with AF2 pLDDT > 90, pAE interaction < 10 Å, and Rosetta binding free energy difference (ddG = ΔΔG) < -40 REU were inputs selected for evaluation by yeast surface display **(left)**. YSD hits were ranked by K_D_ estimate based on sequence counts at each concentration and the 96 designs with estimated K_D_ < 10 nM were elected for recombinant expression in *E. coli*. Of the 96 hits, 4 demonstrated high soluble expression and detectable binding signal (titrated 5x over 6 steps from 5 µM to 1.6 nM in the presence of immobilized NiV F trimer) **(right)**. **(c)** In an SPR competition assay, all 4 binders reduced 1F5 analyte binding to immobilized NiV F trimer **(top)**. These designs also inhibited viral entry with IC_50_ values ranging from 40 to 242 nM. Technical replicates (n=4) at each concentration represented as a mean (dot) and standard deviation (vertical error bar). IC_50_ and standard error of the fit reported in the figure panel. Minibinder A7 demonstrated the most potent *in vitro* neutralizing activity against HIV pseudotyped with the NiV-M F & G glycoproteins **(bottom)**. Minibinder A7 was dubbed F minibinder v1 in efforts to improve affinity and solubility. **(d)** Iterative optimization of minibinder A7 for high affinity binding to NiV F while maintaining soluble expression. **(e)** Cryo-EM structure of F minibinder v3 (green) bound to the prefusion-stabilized NiV F trimer (gray) determined at 2.5 Å resolution **(left)**. Structural overlay of minibinder structure (green) with NiV F design model (yellow) shows agreement, RMSD = 0.3 Å **(top right)**. Interface residues **(bottom right)** show selected interactions.

To identify the most tightly binding designs, yeast surface display was employed as previously described (Cao et al. 2022), sorting against prefusion-stabilized NiV F and HeV F to enrich for cross-reactive binders (**Figure 2b**). The top ninety-six sequences ranked by estimated affinity at the lowest concentration of NiV & HeV F were recombinantly expressed and purified from *E. coli*, and binding was measured by SPR. Four designs bound to NiV F (**Figure 2b**) with K_D_ values ranging from 16-121 nM, competed with antibody 1F5 for binding to NiV F (**Figure 2c**), and inhibited viral entry in a concentration-dependent manner with IC_50_ values ranging from 40-242 nM (**Figure 2c**, **Supplementary Figure 5**, **Supplementary Table 1**). Following affinity maturation using the partial diffusion-ProteinMPNN-AF2 workflow described for the G minibinders (**Figure 2d**), the top hit (F minibinder v2) displayed improved affinity due to off-rate improvements (K_D_ = 1.44 nM , k_off_ = 5.57 × 10^−4^ s^−1^), although evidence of higher order aggregation could explain its slower on-rate (3.85 × 10^5^ M^−1^ • s^−1^). To improve soluble expression without compromising binding potency, the interface of the lead candidate was fixed and the rest of the backbone redesigned with ProteinMPNN with soluble weights. A final round of screening yielded 19/36 soluble and monodisperse binders with the highest binding affinity of 199 pM for NiV F. The most potent minibinder, dubbed F minibinder v3, was tested for cross-reactivity against immobilized NiV F, HeV F, GhV F, and LayV F on SPR alongside earlier versions F minibinder v1 and F minibinder v2 (**Figure 2d**, **Supplementary Figure 6**). For all three F minibinders, binding was observed with NiV F and HeV F, but not for GhV F and LayV F, suggesting that these two distant lineages introduce substitutions on their fusion protein that preclude broader cross-reactivity at this epitope (**Figure 2a**).

We determined a cryoEM structure of the F minibinder v3 in complex with prefusion-stabilized NiV F at 2.5 Å resolution revealing close agreement with the designed model (1.26 Å C_α_-RMSD) when superimposing NiV F (**Figure 2e, Supplementary Figure 7**). Each minibinder buries approximately 825 Å at the interface with Nipah F through contacts formed with a single protomer. Several hydrogen bonds contribute to the strong interactions observed, such as between residues N36_MB_ and A249_NivF_, Q43_MB_ and L285_NivF_, as well as S46_MB_ and T286_NivF_ in the second alpha helix and residues Q52_MB_, D56_MB_ and T59_MB_ with N51_NivF_ as well as Q63_MB_ with T54_NivF_ in the third alpha helix. Based on the F minibinder v3 binding pose, we hypothesize that it would interfere with the conformational changes leading to membrane fusion, explaining its potent neutralizing activity.

### Minibinder oligomerization boosts neutralization potency

As noted above, the high affinity multivalent interactions between EFNB2 and HNV G present a high barrier to neutralization by competitive inhibition. It is possible that the tetravalent NiV G protein, bound by bivalent m102.4, retains unoccupied EFNB2-binding sites that permit host receptor attachment and subsequent viral invasion. To simultaneously address both the affinity and valency hypotheses with high avidity homo-oligomers, we constructed a library of 92 previously characterized cyclic oligomers (ranging from C1-C8) suitable for flexible fusion of the minibinders to the N-or C-terminus via glycine-serine repeat linkers (**Figure 3a**). Each library member contains the sequence of one oligomer and a terminal tandem SNAC tag and 6x-His tag. We hypothesized that highly avid minibinder-oligomer fusions would potentiate viral neutralization over the monomer by increasing complex half-life with the target, enhancing resilience to viral escape (Cao et al. 2020; Hunt et al. 2022; Lee et al. 2026; Ragotte et al. 2025), and reducing the fraction of unbound HNV G protein receptor binding sites.

**Figure 3.**
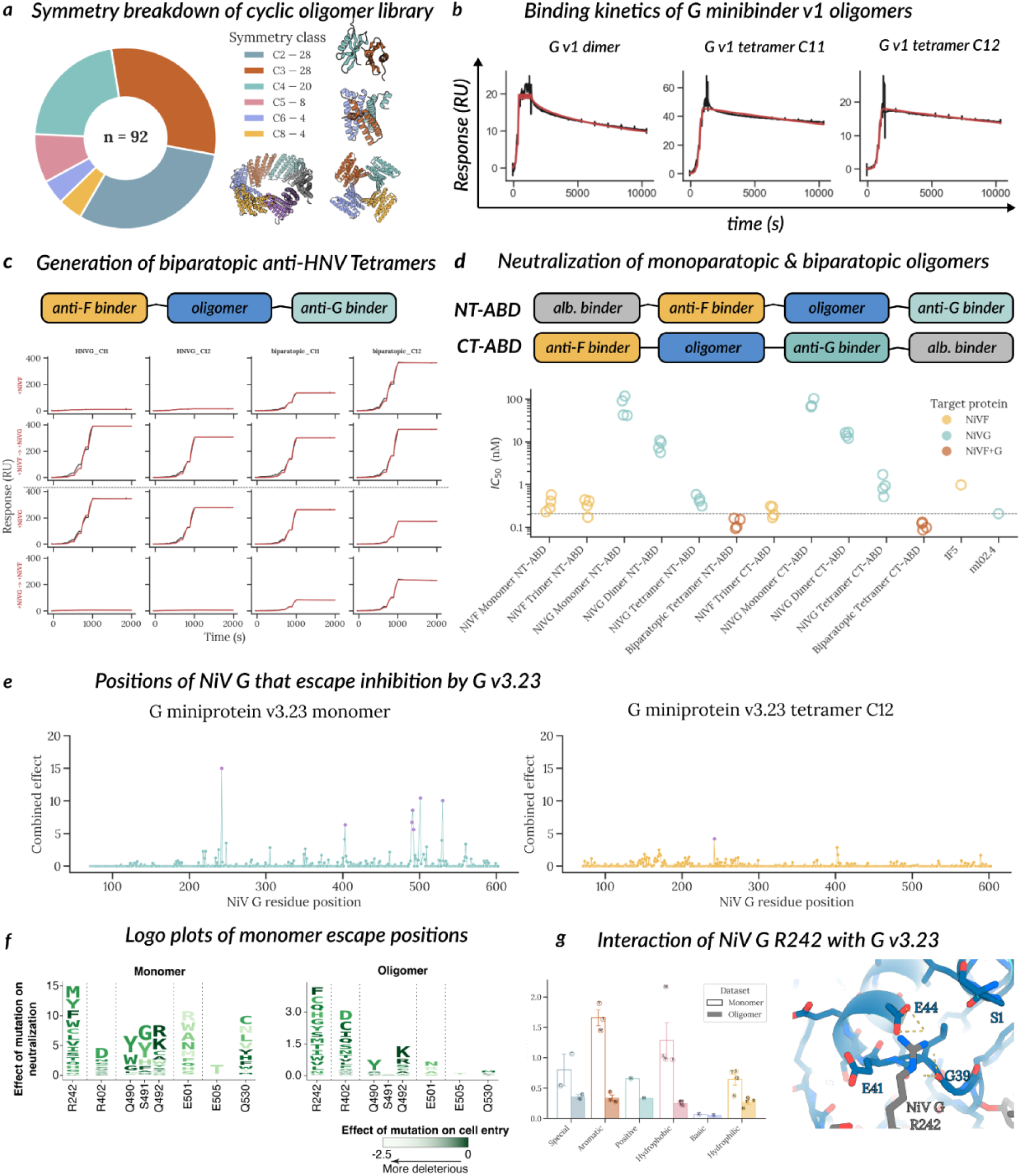
Minibinder oligomerization enhances neutralizing activity and resistance to escape mutations. **(a)** Overview of *de novo* cyclic oligomer library ranging from C2-C8. Flexible GS linkers were used to fuse & display minibinders from the N-or C-termini. **(b)** SPR binding kinetics of the oligomerized G minibinder v1 demonstrate prolonged complex half-life when reacting with immobilized NiV G (compare with kinetics in Fig 1c-middle). **(c)** Evaluation of binding to NiV F & G using monoparatopic and biparatopic tetramers. Monoparatopic tetramers C11 and C12 had C-termini flexibly fused to G minibinder v3, whereas biparatopic tetramers had F minibinder v3 fused to the N-terminus and G minibinder v3 fused to the C-terminus of the tetramer. Immobilized antigen indicated in rows 1 & 3 on the left axis in red (row 1 –NiV F, row 3 NiV G). Secondary “sandwich” antigen binding as analyte indicated in rows 2 & 4 after arrow in red (row 2 – NiV F → NiV G, row 4 – NiV G → NiV F). While monoparatopic tetramers exclusively interacted with NiV G, biparatopic tetramers demonstrated binding when complexed to immobilized NiV G/F, washed, and subsequently exposed to the other NiV glycoprotein (F/G, respectively). **(d)** Overall comparison of neutralization of F & G minibinder v3 monomers & oligomers using a non-replicating, HIV backbone pseudotyped with NiV F & G. Each data point is presented as a biological replicate (n=4, except mAbs where n=1), consisting of the average IC_50_ value obtained from at least two technical replicates. Biological replicates are independently produced batches of pseudovirus and protein. An albumin-binding domain (for extending serum half-life) was included in this study to evaluate the effect of this module on neutralization. While F minibinders demonstrated robust neutralization regardless of oligomerization state, G minibinder demonstrated increased potency with increasing valency (monomer → dimer → tetramer). Ultimately, biparatopic tetramers had the highest potencies of all oligomers and m102.4. **(e)** Total effects of mutations at each site on neutralization as assessed by pseudovirus deep mutational scanning for anti-HNV G minibinders v3.23 monomer (top) and tetramer C12 (bottom). The escape score is defined as log_2_ of the ratio of the frequency of a variant post-selection to the frequency of a mutation pre-selection. Summed escape is the addition of all of the mutation-specific escape scores for a given position. The 6 positions noted to have significantly high escape (p < 0.05) for the monomer were R242, P403, Q490, S491, Q492, and Q530. For the tetramer, only R242 reached statistical significance (see methods), while all other mutation positions regressed to the mean. Logo plots of the statistically significant positions of G minibinder v3.23 monomer are included as part of the inset for the monomer and the tetramer. **(f)** Logo plots of escape positions for the G minibinder v3.23 monomer and for the corresponding oligomer C12. Height of letters indicates the magnitude of decrease in neutralization relative to the unmutated residue. Residues are colored according to their impact on entry in *P. alecto* Ephrin-B3-expressing cells according to the color key. *N.B.*, y-axes are scaled differently for monomer versus tetramer for ease of visualization **(g)** Model-level analysis of R242 demonstrating the involvement of hydrogen-bond acceptors on the minibinder – namely residues E44 (salt-bridge) and G39 (hydrogen bond)), with potential interactions from S1 and E41.

To evaluate the avidity gain resulting from multimerization, we designed 8 oligomeric versions of G minibinder v1 (**Figure 3b**) and found that they bound immobilized NiV G with limited dissociation after 10,000s, confirming the hypothesis that multivalent binding of minibinders to NiV-G prolongs complex half-life. We similarly tested 12 oligomers of the G minibinder v3 (n = 7) along with the G minibinder v3.23 sequence variant (n = 5) spanning C2-C8 geometries by SEC and mass photometry (see Methods). 6/6 of C3+ oligomers had peaks at the expected molecular weight, and 2/6 were monodisperse, indicating that this subset of designs form robust oligomeric assemblies.

To assess developability, G minibinder v3 monomers and oligomers were subjected to stability experiments. First, monomers and oligomers were assessed for thermal stability with circular dichroism spectroscopy. All proteins showed thermal stability up to 75°C and some evidence of partial loss of secondary structure at 85-95°C (**Supplementary Figure 8** & **Supplementary Figure 9**). In an accelerated shelf-life study, monomers and v3 tetramers designated C11 and C12 were incubated for one week in SPR assay buffer (HBS-EP(+)) at temperatures ranging from 4 °C to 65 °C. All proteins retained binding across the temperature range (with marginal loss of signal amplitude at 65°C), demonstrating that retention of folding and binding to the viral target were robust to temperature stress (**Supplementary Figure 10** and **Supplementary Figure 11**). Moreover, the monomers were subjected to overnight incubation in human serum at 37 °C and had binding kinetics assessed with SPR. When compared to a fresh dilution into human serum prior to kinetic evaluation, there was no detectable loss of binding (**Supplementary Figure 12**).

We reasoned that incorporation of a potent and soluble anti-F minibinder on our anti-G oligomers could confer an additional neutralization advantage and superior resistance to escape. With *in vivo* studies in mind, we also incorporated an albumin-binding domain to the N-or C-terminus (NT-ABD vs CT-ABD) to prolong serum half-life. The biparatopic and monoparatopic oligomers were solubly expressed in *E. coli* and robust to overnight room-temperature SNAC cleavage with minimal aggregation (**Supplementary Figure 14**). The biparatopic oligomers were more potent in neutralization than the monoparatopic anti-G oligomers, highlighting the improved neutralization potency with a biparatopic formulation (Figure 3d). The NT-ABD and CT-ABD variants showed similar neutralization IC_50_ values to the corresponding ABD-free designs, suggesting that the ABD does not interfere with binding or neutralization and would allow *in vivo* evaluation via systemic administration (**Figure 3c**, **Figure 3d, Supplementary Figure 13**).

### Lead candidate oligomers reduce HNV escape potential

We sought to prospectively explore the effect of residue substitution on inhibition of viral entry for G minibinder v3.23 (which share identical contact residues with v3) and a tetravalent oligomer version (v3.23 tetramer C12) using a non-replicating, pseudovirus-based deep mutational scanning library of the NiV G glycoprotein, as previously described (Larsen et al. 2025). We titrated the monomer and oligomer against NiVG mutant pseudovirus libraries prior to addition of CHO target cells stably expressing *Pteropus alecto* EFNB3 (bENFNb3). We then used deep sequencing to measure the amount of viral entry into target cells. A non-neutralizing DNA barcode standard was included as a quantification standard.

For the monomer, seven positions were identified with total escape above the mean – R242, P403, Q490, S491, Q492, E501, and Q530 (p < 0.05, Fig. 3a–top, see Methods for statistical analysis), all corresponding to NiV G positions that contact the binding interface (**Figure 3e**). However, with the tetravalent oligomer, we observed that 5/6 of these interface escape mutations regress back to the mean – demonstrating a much higher bar for single-mutation escape (**Figure 3e – bottom**). Similar work on coronaviruses supports the hypothesis that oligomerization of neutralizing moieties mitigates single-mutation escape liabilities and provides a way to maintain antiviral breadth (Hunt et al. 2022; Ragotte et al. 2025; Cao et al. 2020; Lee et al. 2026). For the oligomer, only position NiVG R242 reached statistical significance (escape score = 3.7, p = 0.005); this is the mutation that conferred the highest escape potential for the monomer (escape score = 13.6, p < 1e-16, **Figure 3e-g)**. However, for the oligomer, all substitutions at R242 had reduced escape scores compared to those of the monomer, and all detected non-basic residues performed similarly (< 0.5) suggesting that oligomerization dramatically reduces the biochemical impact of substitutions on reducing viral entry. R242 interacts with hydrogen-bond acceptors (S1 side-chain hydroxyl, G39 backbone carbonyl O, E41 side-chain hydroxyl, and E44 side-chain hydroxyl; G39 and E41 derived from the Ephrin G-H loop) (**Figure 3g**). Redesign of the v3.1 and v3.23 S1 and E44 could further reduce escape potential. These data show that oligomerization of a G minibinder markedly dampens susceptibility to escape mutations, a favorable property for antiviral inhibitors.

Although Nipah and Hendra viruses have limited genetic divergence, the recent discovery of a new Hendra virus genotype, designated HeV-g2, underscored the continued risk of emergence of divergent henipaviruses (J. Wang et al. 2021; Annand et al. 2022). HeV-g2 F and G respectively share 95.60% and 92.85% amino acid sequence identity with prototypic HeV F and G (Z. Wang, Dang, et al. 2022). Based on the strict conservation of the epitopes targeted by the lead F and G minibinders described here between prototypic HeV and HeV-g2, we anticipate that these F and G minibinders will neutralize viral genotypes as divergent as HeV-g2.

## Discussion

F and G are the primary targets of neutralizing antibodies which correlate with protection in animals infected with NiV or HeV (Bossart et al. 2009; Zeitlin et al. 2024; Geisbert et al. 2014; Mire et al. 2020; Dang et al. 2021, 2019; Xu et al. 2013; Avanzato et al. 2019; Byrne et al. 2023). The G-directed, NiV and HeV cross-reactive m102.4 mAb has been administered on a compassionate use basis to 15 individuals with high-risk exposure and completed a phase 1 clinical trial. The humanized version of the F-targeting 1F5 mAb outperformed m102.4 when administered post-infection in a non-human primate challenge model with NiV-B and is about to enter clinical trials (Zeitlin et al. 2024). Owing to the requirement for two distinct glycoproteins for HNV entry into cells, recent designs have targeted both F and G through the usage of bispecific antibodies and cocktail of mAbs (Isaacs, Nieto, Zhang, Modhiran, Barr, Thakur, Low, Parry, Barnes, Jara, Himelreichs, Yao, Deride, Barthou-Gatica, Salinas-Rebolledo, Ehrenfeld, et al. 2025; Guzmán-Solís et al. 2026). Based on these encouraging results, we set out to generate bi-specific constructs that bind to F and G through small designed binding domains; such miniprotein binders are an emerging and promising therapeutic modality due to their smaller size, high stability, lower cost of expression, and comparable potencies to neutralizing monoclonal antibodies (Ragotte et al. 2025; Case et al. 2021).

We describe here the modular design of anti-henipaviral miniproteins with best-in-class potencies and favorable drug-like properties (solubility, thermostability, manufacturability). We show that the miniproteins can be readily oligomerized to increase avidity and neutralization potency and can be conjugated to albumin-binding domains to increase serum half-life in a sequence-efficient manner: our most potent design is ∼40% of the molecular weight of an antibody and has the greatest potency of any HNV inhibitor described to date. As illustrated by the excellent affinities and solubilities of the best designs described here, our partial diffusion-soluble MPNN approach allows sequence optimization for both affinity and solubility and should be readily applicable to other designed binders. The favorable biochemical properties of these inhibitors will allow exploration of different routes of administration, and they represent strong preclinical candidates for further evaluation. Both F and G minibinders potently inhibit NiV and HeV F/G-mediated entry into cells and the lead NiV F minibinder achieves greater neutralization potency than the 1F5 mAb. Consequently, these minibinders are expected to be protective *in vivo*. Using deep mutational scanning, we show that oligomerization of the G minibinder heightens the barrier for emergence of neutralization escape mutants. We therefore anticipate that targeting of F and G simultaneously with our biparatopic minibinders would increase resistance to emergence of escape mutations due to the requirement of a minimum of two mutations to abrogate recognition by each minibinder module, making it a promising therapeutic candidate.

## Acknowledgements

This work was supported in part by the NIH/NIAID under grants 1U19AI181881 to C.C.B., J.D.B., D.V., and D.B., DP1AI158186 and 75N93022C00036 to D.V., an Investigators in the Pathogenesis of Infectious Disease Awards from the Burroughs Wellcome Fund to D.V., the University of Washington Arnold and Mabel Beckman Cryo-EM Center. M.A. and C.C.B. were supported by NIH grants AI181930 and AI142764. J.D.B., D.V. and D.B. are investigators of the Howard Hughes Medical Institute, and D.V. holds the Hans Neurath Endowed Chair in Biochemistry at the University of Washington.

## Contributions

JS, DV, and DB designed the project. JS, RG, KS, YJP, ZW, BL, MA, and DV designed experiments. JS designed and characterized the minibinders and oligomers and evaluated binding to recombinant antigens. KS carried out pseudovirus neutralization assays and MA carried out rCedV neutralization assays. RG, YJP, and ZW performed sample vitrification, cryoEM data collection and processing. RG, YJP built and refined atomic models with assistance from DV. RG designed constructs. RG, ZW, CS, and JB expressed recombinant glycoproteins. BL carried out pseudovirus deep mutational scanning and data analysis. JS, RG, KS, DV, and DB analyzed the data. JS, RG and DV wrote the manuscript with input from all co-authors. CB, JB, DV, and DB acquired funding and supervised the project.

## Declaration of interests

MA and CCB are United States federal employees and co-inventors on United States and foreign patents pertaining to Cedar Virus and Methods of Use, whose assignees are the United States as represented by the Henry M. Jackson Foundation for the Advancement of Military Medicine, Inc. (Bethesda, MD, USA). All other authors declare no conflict of interest.

## Disclaimer

The views expressed in the manuscript are solely those of the authors, and they do not represent official views or opinions of the Uniformed Services University, the United States Department of War or the Henry M. Jackson Foundation for the Advancement of Military Medicine, Inc. Several of the authors are U.S. Government employees. This work was prepared as part of their official duties. Title 17 U.S.C. § 105 provides that ‘Copyright protection under this title is not available for any work of the United States Government.’ Title 17 U.S.C. §101 defines U.S. Government work as work prepared by a military service member or employee of the U.S. Government as part of that person’s official duties.

## Materials & Methods

### Computational design of G minibinders

To generate scaffolds for the function NiV G-binding motif, we provided a motif of Ephrin B2 (G-H loop and preceding β-strand, residues 109-128) and a truncated version of NiV G that maintains all interface contacts from PDB 2VSM to reduce total run time. To ensure that appropriate scaffold diversity could be generated without restricting the overall position of the fixed EFNB2 motif, RFdiffusion was permitted to sample from a range of 15-60 residues on either side of the motif, allowing a maximum backbone length range of 55-100 residues (including the EFNB2 motif). A total of 18,933 backbones were generated, which then served as input for sequence design.

To assign amino sequences to each backbone, ProteinMPNN was alternated with FastRelax as previously described ((Watson et al. 2023), Methods). During sequence design, we fixed the residue identity of the scaffolded EFNB2 motif to design a sequence around the motif that would scaffold the EFNB2 interface without compromising its structure and function. A total of 37,873 design models with unique sequences were used as input to AlphaFold2 (AF2, (Jumper et al. 2021) for single sequence folding prediction without multiple sequence alignment.

### Computational design of F minibinders

NiV F (PDB code: 6TYS) was elected as a target for minibinder generation. Hotspots were based on hydrophobic patches, favorable secondary structural motifs (like exposed β-strands), and reasonable distance from glycans. These hotspots were used to guide RFDiffusion trajectories for sequences ranging from 55-100 residues in length – a total of 54,873 backbones were generated. Sequence design with ProteinMPNN (without fixing any motifs) and structural prediction with AF2 were performed as described with the design of the G minibinders. Additionally, Rosetta energies were scored on relaxed AF2 structures, and 1601 candidates with favorable folding and complexation score predictions (pLDDT ≥ 85, pAE_interaction < 10, and Rosetta ΔΔG score < -40) were selected for yeast surface display.

### Computational optimization of F G G minibinders

v1 hits identified with SPR were subjected to another round of partial diffusion using RFDiffusion followed by sequence assignment with ProteinMPNN for the purpose of identifying structural and sequence variants with higher binding potency. In the case of G minibinders, the G-H loop was fixed during partial diffusion. The same folding and filtering schema elucidated in the respective computational design sections above was employed to identify hits, which were then screened again for soluble recombinant expression and binding at 96-well scale (see 96-well, 4-mL protein purification and size-exclusion chromatography section below in Methods).

To generate v3 designs, v2 hits were then subjected to sequence redesign with solubleMPNN (ProteinMPNN with published soluble weights) while fixing the interface residues. These designs were then folded and filtered *in silico* using the same metrics as before, and then screened at 96-well scale for SEC monodispersity and on-target peak as well as higher affinity than the v2 parent design prior to clonal isolation and 50-mL scale up.

### Yeast surface display and data analysis of designed miniproteins

Yeast surface display – included adapter design, transformation, and sorting – was employed as described in (Cao et al. 2022). In brief, amplified DNA oligos were transformed into chemically competent *S. cerevisiae* EBY100, and cells were recovered in C-Trp-Ura medium supplemented with 2% (w/v) glucose (CTUG) at 30°C 225rpm overnight. Cells were induced for expression with SGCAA media supplemented with 0.2% (w/v) glucose for 12-16h with shaking at 30°C 225rpm. Cells were then washed with 1x PBS + 1% Bovine Serum Albumin (PBSF). For the first phase of sorting, cells were stained with anti-c-Myc fluorescein isothiocyanate (FITC, Miltenyi Biotech) and sorted for expression. The second sort (avidity sort) was stained with a pre-incubated mix of 1 µM biotinylated NiV F + 1 µM protomer streptavidin–phycoerythrin (SAPE, ThermoFisher) + FITC. All subsequent sorts were performed by incubating induced cells for 1h at 4°C with biotinylated NiV or HeV F glycoprotein, followed by staining with 1:1000 SAPE & 1:1000 FITC at 4°C for 10 minutes. Sort 3 used 1 µM biotinylated HeV F trimer to select for cross-reactive minibinders. Sort 4 titrated biotinylated NiV F trimer (1 µM, 100 nM, 10 nM, 1 nM, 100 pM). K_D_ upper bound and lower bound was estimated by comparing NGS counts at different antigen concentrations, as described previously (Cao et al. 2022).

### Golden gate assembly and chemical transformation

As previously described (Watson et al. 2023), DNA synthesized as E-blocks from IDT encoding the designed protein of interest were cloned into pET-29b(+) vectors with C-terminal tandem SNAC and 6x-His tags modified with BsaI cut sites, leaving compatible sticky ends (5’ –AGGA–gene of interest–TTCC – SNAC – 6x-His – 3’) following vendor instructions for BsaI-HF (New England Biolabs). Plasmid was transformed without cleanup into chemically competent BL21-DE3 (New England Biolabs), and cells were recovered for 1 hour at 37 °C in SOC Outgrowth Medium (New England Biolabs). In the 96-well format, transformants were used to inoculate 4 x 1 mL plates of auto-induction media (Watson et al. 2023), and a 1 mL plates with LB + 50 µg/mL kanamycin sulfate was prepared for glycerol stocks after overnight growth at 37 °C with shaking at 1000 rpm. Otherwise, transformants were spread on plates with LB + 100 µg/mL kanamycin sulfate, and colonies were sequenced. Colonies were then used to inoculate 50 mL cultures in 250-mL baffled flasks for 20h growth at 37 °C with shaking at 250 rpm.

### HNV protein and antibody construct design

Nipah F sequence (Uniprot: Q9IH63 ) residues 1-487 was cloned into a CMVR construct with a C-terminal GCN4 trimerization sequence and 8XHis tag, along with disulfide mutations N100C-A119C and Y97C-G131C, A156P, S470V, Y473E, and A477V (Langedijk et al. 2024). The Nipah G head construct includes Niv G residues 176-601 fused to a N terminal 6xHis tag into pOPING vector with a Mu-phosphatase signal peptide. The HENV-32 Fab sequences were retrieved from the crystal structure PDB 6VY4, and cloned into a pcDNA3.1(+) vector with a signal sequence (MPMGSLQPLATLYLLGMLVASVLA) at the N-termini and Strep tag II at the C-terminus for the heavy chain. A consensus sequence of IGHG1 (P01857) was used to yield a full-length heavy chain sequence. All constructs were synthesized by Genscript.

### 50-mL E. coli expression, purification, and SEC of designed proteins

Proteins were purified as described previously (Watson et al. 2023). Gene fragments encoding the 22 designs (all ∼10 kDa) were cloned into a pET-28b(+)-based expression plasmid, expressed in BL21(DE3) *E. coli,* and expressed at 50mL scale (37°C, 225 rpm, 20h) with a C-terminal tandem SNAC tag and 6x-His tag as previously described. Cell pellets were harvested with centrifugation (4000xg, 10 min, 25°C), resuspended in lysis buffer (25 mM Tris HCl pH 8, 300 mM NaCl, 40 mM Imidazole, 1 mM DNase I, 10 µg /mL lysozyme), and and cells were lysed via sonication (5 min, 85% amplitude, 15s on/off cycles). Centrifugation was used to clarify lysate (14,000 x *g*, 20 min., 4°C), prior to its addition to 1-mL Ni-NTA columns pre-equilibrated with wash buffer (25 mM Tris HCl pH 8, 300 mM NaCl, pH 8). Resin was washed with 5 column volumes of wash buffer, and proteins were eluted in 1.6 mL elution buffer (50 mM Tris HCl pH 8, 300 mM NaCl). Samples of 1 mL were injected onto a Superdex-75 Increase 10/300 GL (Cytiva) or a Superdex-200 Increase 10/300 GL (Cytiva) for monomers and oligomers, respectively, and absorbance at 280 nm was used to track protein elution.

For designs tested *in vitro* or for cryo-EM studies, SNAC tags were cleaved on Ni-NTA resin by washing with 5 CVs (5 mL) of cleavage buffer (100 mM CHES, 100 mM Acetone oxime, 100 mM NaCl, 500 mM GnCl, pH 8.6) prior to overnight incubation at room temperature in 40 mL SNAC cleavage buffer + 2 mM Ni(II)Cl_2,_ or Ni(II)SO_4_. The eluate was collected from a filtered gravity column and was then concentrated with a 3 kDa (for monomers) or 10 kDa (for oligomers) spin concentrator (Amicon Ultra 4/15 mL centrifugal devices with Ultracel®) prior to sizing.

### 25-mL Expi293 expression, purification, and SEC of HNV proteins and mAbs

Purification of Nipah F, G head, and HENV-32 mAb for cryoEM sample preparation Nipah F ectodomain and G head were produced in 25mL Expi293F cells grown in suspension using Expi293 Expression Medium (Thermo Fisher) at 37°C in a humidified 8% CO2 incubator rotating at 130 rpm. The cultures were transfected using ExpiFectamine™ 293 Transfection Kit for Expi293 cells (Thermo Fisher) with cells grown to a density of 3 million cells per mL and cultivated for 5 days for Nipah F and 6 days for Nipah G. The supernatants were harvested, and proteins were purified from clarified supernatants using a 1 mL HisTrap HP for Nipah F and 1mL Cobalt affinity column (Takara) for Nipah G. Nipah G head was buffer exchanged, concentrated and flash frozen in TBS (50 mM Tris, 150 mM NaCl and 10 mM EDTA at pH 8.0). SDS-PAGE was run to check purity. Nipah F was further purified via size exclusion chromatography on a Superdex200 10/300 GL (Cytiva), and stored in 25 mM Tris, 150 mM NaCl at pH 8.0.

HENV-32 mAb heavy and light chains were transfected at a 1:1 ratio in Expi293 cells and cultivated for 6 days. The supernatants were harvested, and proteins were purified from clarified supernatants using a 1 mL StrepTrap^TM^ HP column (Sigma), buffer exchanged, and concentrated and flash frozen in TBS. SDS-PAGE was run to check purity. LysC digestion was performed to generate the Fab.

1F5 mAb heavy and light chains were transfected at a 1:1 ratio in ExpiCHO cells and cultivated for 5 days. The supernatant was harvested, and proteins were purified from clarified supernatants using a 1 mL HisTrap column (Cytiva), buffer exchanged, concentrated, and flash frozen in TBS. SDS-PAGE was run to check purity.

Human mAb m102.4 is a cross-reactive and neutralizing NiV and HeV G glycoprotein specific IgG1 (Zhu Z, et al., PMCID: PMC7199872; doi: 10.1086/528801). m102.4 (lot: PURSR2-02) was produced and purified, involving protein A affinity chromatography, from master cell bank CHO-K1 m102.4 IgG1 by Laureate Biopharma, Princeton, NJ, under SRI Project P17545.294 (SRI International, Menio Park, CA), and supported by DMID, NIAID, NIH Services for Preclinical Development of Therapeutic Agents Contract.

### Octet biolayer interferometry

NiV G (prepared as described in (Cao et al. 2022; E. Y. Wang et al. 2022)) was chemically conjugated as ligand to tips from an Amine coupling kit (Sartorius) according to manufacturer instructions in acetate buffer pH 5. Binders were diluted serially in octet buffer (HBS-EP (+), pH 7.4, Cytiva) as analyte 2x from 1µM in 6 wells, with two wells reserved as no-analyte and no-ligand controls. After establishing a baseline in octet buffer, ligand was exposed to binder for association and then placed into octet buffer for dissociation. The same setup was repeated for successful binders with binder immobilized as ligand, and NiV G titrated as analyte.

### 96-well, 4-mL purification and size-exclusion chromatography (SEC) of designed proteins

Proteins were purified similarly (described in (Watson et al. 2023)) to the 50-mL protocol with a few exceptions. After pellet isolation, pellets were lysed with B-PER™ Bacterial Protein Extraction Reagent (Thermo Scientific) at 37 °C for 15 min at 250 rpm. Plate-based Ni-NTA pulldowns used 50 µL Ni-NTA resin, and proteins were eluted in 100 µL elution buffer, spun through a 0.22 µm filter plate (Pall – AcroPrep Advance 96-well Filter Plates -2 mL, 0.2 µm wwPTFE membrane). Sizing was performed on a Superdex-75 Increase 5/150 GL (Cytiva) or a Superdex-200 Increase 5/150 GL (Cytiva) for monomers and oligomers, respectively.

### Surface plasmon resonance (SPR) kinetic measurements

NiV & HeV G (prepared as described in (Cao et al. 2022; E. Y. Wang et al. 2022)) was immobilized on a CM5 SPR chip (Cytiva) in acetate buffer, pH 5, and binder plates were prepared with 6 x 5-fold serial dilutions in HBS-EP (+), pH 7.4 (Cytiva) for a single-cycle kinetics experiment, as described previously ((Watson et al. 2023)). To assay reactivity to EphB4, we immobilized Fc-tagged EphB4 (AcroBiosystems) to a protein A chip (Cytiva) for single-cycle kinetics experiments. Analyte proteins were titrated from 5 µM to 1.6 nM over 6 x 5-fold serial dilutions.

### NiV G pseudovirus deep mutational scanning

Deep mutational scanning Nipah virus G deep mutational scanning libraries were produced as previously described (Larsen et al. 2025). These experiments were performed with nonreplicative lentiviral-based pseudoviruses that can only undergo a single round of infection and do not encode any viral genes other than the Nipah virus G protein. To ensure we do not describe information about human-specific adaptations that could be misused, we performed all experiments in CHO cells that stably express EFNB3 from a natural henipavirus host, *Pteropus alecto*. To measure how Nipah virus G mutations affect neutralization by protein minibinders, we incubated the libraries for 1 hour at 37°C with protein dilutions that neutralize ∼50 and 95% of the library viruses. Library mixtures were used to infect CHO-bEFNB3 cells. Twelve hours post-infection, lentiviral DNA templates were recovered, followed by PCR amplification of 16 nt barcodes and Illumina sequenced. All experiments were performed in duplicate with independent pseudovirus libraries. Data were analyzed with dms-vep-pipeline-3 (Dadonaite et al. 2023) and the effects of mutations on neutralization were calculated with a biophysical model in polyclonal (https://jbloomlab.github.io/polyclonal/) (Y. Liu et al. 2022). The analysis pipeline and filtered results are available on GitHub: https://github.com/dms-vep/Nipah_Malaysia_RBP_Minibinder_DMS.

We identified mutations that reduced neutralization by the minibinders using a previously described pseudovirus deep mutational scanning approach involving libraries of pseudotyped lentiviral particles encoding the Nipah virus G (also known as RBP) (Larsen et al. 2025). These pseudoviruses encode no viral proteins other than NiV G, and so are not fully infectious agents capable of causing disease, making them a safe way to study the effect of mutations to Nipah G at biosafety level 2. In brief, anti-NiV G minibinders (v3.23 monomer and v3.23 oligomer C12) were incubated at different concentrations corresponding to IC50 and IC95 with ∼1 × 10^6^ TU of pseudovirus library followed by addition to CHO-bEFNB3 cells. After a 12-16h incubation, DNA was extracted and barcodes were quantified by Illumina next-generation sequencing.

The deep mutational scanning data were filtered & processed according to the criteria described in (Larsen et al. 2025). To identify positions of escape, a standardized z-score was computed for each G-protein residue position after summing the escape of all variants at each position. One-sided p-values were derived assuming asymptotic normality of the standardized z-statistic. Benjamini–Hochberg false discovery rate (FDR) correction was implemented to control for multiple comparisons across residues, with significance defined as FDR < 0.05. Positions with insufficient data for statistical analysis were excluded from testing. All data, code, and results from the deep mutational scanning experiments can be found here (https://github.com/dms-vep/Nipah_Malaysia_RBP_Minibinder_DMS).

### Fluorescence reduction neutralization test (FRNT)

Recombinant Cedar viruschimeras (rCedV-NiV-B-GFP and rCedV-HeV-GFP) were rescued and handled following laboratory manipulation guidelines and standard operating procedures under BSL-2 conditions that were developed, reviewed, and approved by the Uniformed Services University, Institutional Biosafety Committee in accordance with NIH guidelines (Laing et al. 2019, 2018). Replication-competent recombinant CedV (rCedV) chimeric viruses were generated to display the G and F proteins of NiV-B and HeV and to encode a GFP reporter, as described previously (Amaya et al. 2023). Vero 76 cells were seeded at a density of 2×10^4^ cells/well in black-walled clear bottom 96-well plates (Corning Life Sciences; Corning, NY, USA) and incubated for 24 hours at 37 °C, 5% CO2. Minibinders were serially diluted 3-fold such that an initial concentration of 1 µg/mL was used for the 7-point dose-response curve. An equal volume of DMEM-10 containing either rCedV-NiV-B-GFP or rCedV-HeV-GFP was added to each dilution for a final MOI of 0.05 and incubated for 2 hours at 37 °C, 5% CO2. Each of the virus-minibinder mixtures (90 µL/well) was added to the pre-seeded Vero 76 cells in duplicate and incubated for an additional 24 hours at 37 °C, 5% CO2. The virus-minibinder supernatants were removed, and the plates were fixed with 4% Formaldehyde in 1X PBS for 20 min at room temperature. The plates were then washed 3 times with diH2O, and the plates were imaged using a CTL S6 analyzer (Cellular Technology Limited; Shaker Heights, OH, USA). Fluorescent foci were counted using the CTL Basic Count software. Neutralization percent (%) was calculated based on fluorescent foci for each virus without minibinder. These data represent mean ± standard deviation from two independent experiments, each performed in duplicate. The 50% inhibitory concentration (IC_50_) was determined as the minibinder concentration at which there was a 50% reduction in fluorescent foci versus untreated control wells. The IC_50_ values were calculated by non-linear regression curve fitting with a variable slope using GraphPad Prism 10 (GraphPad Software Inc., San Diego, CA, USA). The limit of detection for this assay was 50 fluorescent foci.

### Cryo-electron microscopy data collection, processing and model building

The minibinder v3 bound NiV G head and HENV32 Fab complex was prepared by mixing a 1:1.2:5 molar ratio of the HENV32 Fab, NiV G head and minibinder v3 before incubation for one hour on ice. 3µL of 1.2 mg/ml complex was applied onto freshly glow discharged R 2/2 UltrAuFoil grids (Russo and Passmore 2014) prior to plunge freezing using a vitrobot MarkIV (ThermoFisher Scientific) with a blot force of 0 and 4.5 sec blot time at 100% humidity and 22°C. And also 3 µL of 1.2 mg/mL complex with 3mM 3-[(3-Cholamidopropyl)dimethylammonio]-2-hydroxy-1-propanesulfonate (CHAPSO) was added to the glow discharged side of R 2/2 UltrAuFoil grids (Russo and Passmore 2014) and 1µL was added to the back side before plunging into liquid ethane using a GP2 (Leica) with 6 sec blot time. The data were acquired using an FEI Titan Krios transmission electron microscope operated at 300 kV and equipped with a Gatan K3 direct detector and Gatan Quantum GIF energy filter, operated in zero-loss mode with a slit width of 20 eV. Automated data collection was carried out using Leginon (Suloway et al. 2005) at a nominal magnification of 105,000x with a pixel size of 0.835 Å. The dose rate was adjusted to 10.5 counts/pixel/s, and each movie was acquired in counting mode fractionated in 100 frames of 40 ms. A total of 7,109 movies were collected with a defocus range between -0.2 and -3 μm and stage. Movie frame alignment, estimation of the microscope contrast-transfer function parameters, particle picking, and extraction were carried out using Warp (Tegunov and Cramer 2019). Particles were extracted with a box size of 192 pixels with a pixel size of 1.67Å. The reference-free 2D classification was performed using cryoSPARC (Punjani et al. 2017) to select well-defined particle images. Initial model generation was carried out using ab-initio reconstruction in cryoSPARC (Punjani et al. 2017) and the resulting maps were used as references for heterogeneous 3D refinement. Particles belonging to classes with the best resolved minibinder-bound NiV G head complex density were selected. To further improve the data, the Topaz (Bepler et al. 2019) model was trained on Warp-picked particle sets belonging to the best classes after 2D classification and particles picked using Topaz were extracted and subjected to 2D classification and heterogenous 3D refinements. The two particle sets from the Warp and Topaz picking strategies were merged and duplicate particles were removed using a minimum distance cutoff of 90 Å. The 3D refinement was carried out using non-uniform refinement with per particle defocus in cryoSPARC (Punjani et al. 2020). The dataset was transferred from cryoSPARC to Relion using the pyem program package (ref: DOI 10.5281/zenodo.3576629) and particle images were subjected to the Bayesian polishing procedure implemented in Relion (Zivanov et al. 2018) during which particles were re-extracted with a box size of 384 pixels and a pixel size of 0.835 Å. After polishing, the reference-free 2D classification was performed using cryoSPARC. Particles belonging to the best class were selected. The final 3D refinements were carried out using non-uniform refinement along with per-particle defocus refinement in cryoSPARC (Punjani et al. 2020) to yield the global reconstruction at 2.77 Å resolution comprising 811,703 particles. To improve the density of the minibinder and NiV G head, the final local refinement (Punjani et al. 2017) was carried out using cryoSPARC with a soft mask comprising variable domain of HENV32 Fab, minibinder v3 and NiV G head resulting in a 2.7 Å resolution reconstruction. Reported resolutions are based on the gold-standard Fourier shell correlation (FSC) of 0.143 criterion and Fourier shell correlation curves were corrected for the effects of soft masking by high-resolution noise substitution (S. Chen et al. 2013; Rosenthal and Henderson 2003). Local resolution estimation was carried out using cryoSPARC. Map sharpening in cryoSPARC was performed to assist model building. UCSF Chimera (Pettersen et al. 2004), Coot (Emsley et al. 2010), and Phenix (Liebschner et al. 2019) were used to fit, build, and refine the model using the sharpened and unsharpened cryo-EM maps. Validation used Phenix, Molprobity (V. B. Chen et al. 2010), EMRinger (Barad et al. 2015) and Privateer (Agirre et al. 2015).

The minibinder v3 bound Nipah F head complex was prepared by mixing a 1:5 molar ratio of Nipah F:minibinder before incubation for 1h on ice. 3 uL of 0.1 mg/mL complex was applied onto freshly glow discharged thin carbon grids prior to plunge freezing using a vitrobot MarkIV with a blot force of -1, 2.5s blot time, and 20s wait time at 100% humidity and 22°C. Automated data collection was carried out using SerialEM (Mastronarde 2005) at a nominal magnification of 105,000x with a pixel size of 0.829 Å. A total of 11,307 micrographs were collected with a defocus range between -0.6 and -1.6 μm and stage tilt angle of 0°. Movie frame alignment, estimation of the microscope contrast-transfer function parameters, particle picking, and extraction were carried out using cryoSPARC Live (Punjani 2020), and all downstream processing steps were done in cryoSPARC. Particles were extracted with a box size of 288 pixels and downsampled by a factor of 2. One round of reference-free 2D classification was performed to select well-defined particle images. Initial model generation was performed using ab-initio reconstruction and the resulting map was used for homogeneous and three rounds of heterogeneous refinement. Re-extracted particles were used for subsequent non-uniform refinement with per-particle defocus. Reference-based motion correction and final non-uniform refinement with per-particle defocus was performed to obtain the final reconstruction at 2.5Å. Reported resolution is based on the gold-standard Fourier shell correlation (FSC) of 0.143 criterion. See also Supplementary Figures 4 and 8. ModelAngelo (Jamali et al. 2024) was used for initial model building with sequences of the minibinder and Nipah F ectodomain. The structure was then manually rebuilt using Coot (Emsley et al. 2010) and refined using Phenix (Liebschner et al. 2019). Validation used Molprobity (V. B. Chen et al. 2010), Phenix, and Privateer (Agirre et al. 2015).

### Pseudotyped Virus Production

A single-cycle (replication-incompetent) HIV-1 envelope-deficient backbone vector containing a luciferase reporter was pseudotyped with Nipah F (fusion, Genbank ID: NP_112026.1) and G (attachment, Genbank ID: NP_112027.1) Malaysia strain glycoproteins. Briefly, HEK293T/17 (ATCC, CRL-11268) were seeded (15-20million cells) in poly-D-lysine-coated, T175 flasks in DMEM (ThermoFisher, Gibco 11-965-118) supplemented with 10% FBS (Cytiva, Hyclone SH3039603HI) and incubated overnight at 37°C +5% CO2. The following day, one tube containing 1.5mLs of Opti-MEM (ThermoFisher, Gibco 31985088), 15ug pHage-CMV-Luc2-IRES-ZsGreen-W, 3.3ug pRC-CMV-ReV1b, 3.3ug HDM-tat1b, 3.3ug HDM-Hgpm2, 5.1ug Nipah full-length F and 5.1ug Nipah full-length G and a second tube of 1.5mL Opti-MEM containing 87.75uL of Lipofectamine 2000 (ThermoFisher, Invitrogen 11668500) were combined and incubated at room temperature for 15 minutes. The transfection mixture was subsequently added to the T175 plates containing washed cells with a volume of 20mL of DMEM supplemented with 10% FBS. Plates are then returned to incubate at 37°C +5% CO2 overnight. After ∼24hrs, supplement plate(s) with 200uL sodium butyrate (from 500mM stock prepared in filtered MilliQ water) to achieve a final concentration of 5mM in 20mL of media. Return to the incubator overnight. After ∼24hrs, collect the supernatant from each flask, concentrate virus 10x using a 30kDa cut-off membrane (Millipore Sigma, Amicon UFC903024). Pseudotypes are then aliquoted and frozen at -80°C until use.

### Pseudovirus Neutralization Assay

For Nipah-HIV pseudovirus neutralization assays, HEK293T/17 (ATCC, CRL-11268) were seeded into 10cm dishes containing DMEM (ThermoFisher, Gibco 11-965-118) supplemented with 10% FBS (Cytiva, Hyclone SH3039603HI) and incubated overnight at 37°C +5% CO2. The following day, when cells reached 90% confluency, they were gently washed with DMEM 3x. Next, 1.5mL Opti-MEM (ThermoFisher, Gibco 31985088) containing 8ug human Ephrin-B2 full-length plasmid and a second tube of 1.5mL Opti-MEM containing 30uL of Lipofectamine 2000 (ThermoFisher, Invitrogen 11668500) were combined and incubated at room temperature for 15 minutes. The transfection mixture was subsequently added to the 10cm plates containing washed cells with a volume of 7mL of DMEM supplemented with 10% FBS. After 3-5hrs, cells were detached, counted and reseeded at ∼40,000 cells per well in poly-lysine-coated 96 well plates. Cell plates were returned to incubate overnight at 37°C +5% CO2. The next day, a half-area 96-well plate was prepared with a 1:3 serial dilution of sera in DMEM in 22 uL final volume. 22 uL of pseudovirus was then added to each well and incubated at room temperature for 30–45 min. The media was removed from transfected HEK293T cells and 40 uL of the sera/pseudovirus mixture was transferred to the cells and incubated for 2 h at 37°C +5% CO2 before adding 40 uL of 20% FBS and 2% PenStrep (200 I.U./mL penicillin and 200 mg streptomycin) containing DMEM. Following 18–22 h incubation, 40 uL of One-GloEX (Promega E8150) was added to the cells placed on a shaker and incubated in the dark for 5 min. Plates were then read on an Agilent BioTek Neo2 plate reader. Relative luciferase units were plotted and normalized in Prism (GraphPad) using a zero value of cells alone and a 100% value of 1:2 virus alone. Nonlinear regression of log(inhibitor) vs. normalized response was used to determine ID50 values from curve fits. At least two biological replicates with two distinct batches of pseudovirus were conducted for each minibinder sample.

## Supplementary Figures

**Supplementary Figure 1.**
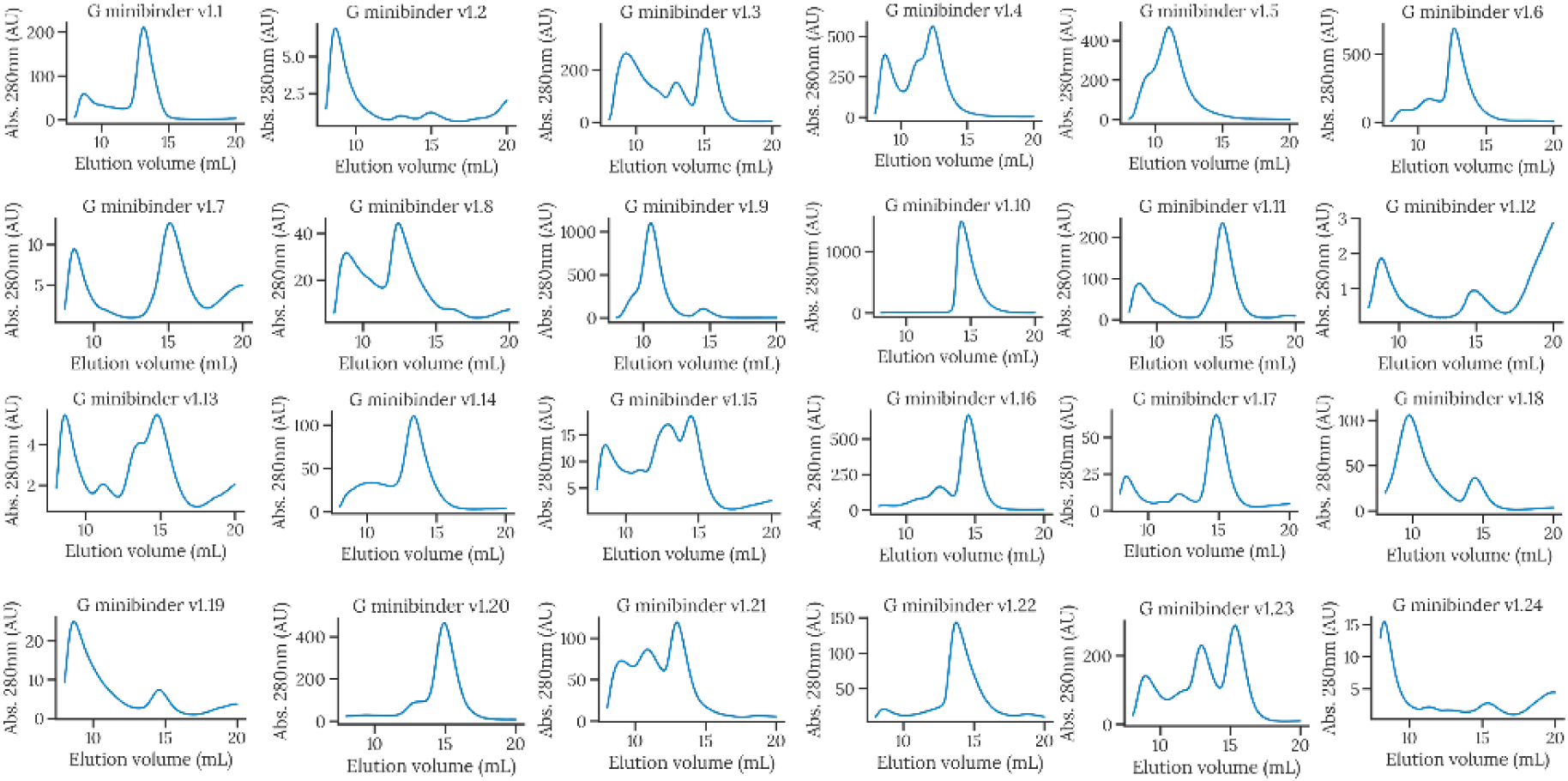
SEC traces (S75 Increase 10/300 GL) curves of G minibinder v1 design series. G minibinder v1.10 identified via this screen became G minibinder v1 for subsequent testing and optimization.

**Supplementary Figure 2.**
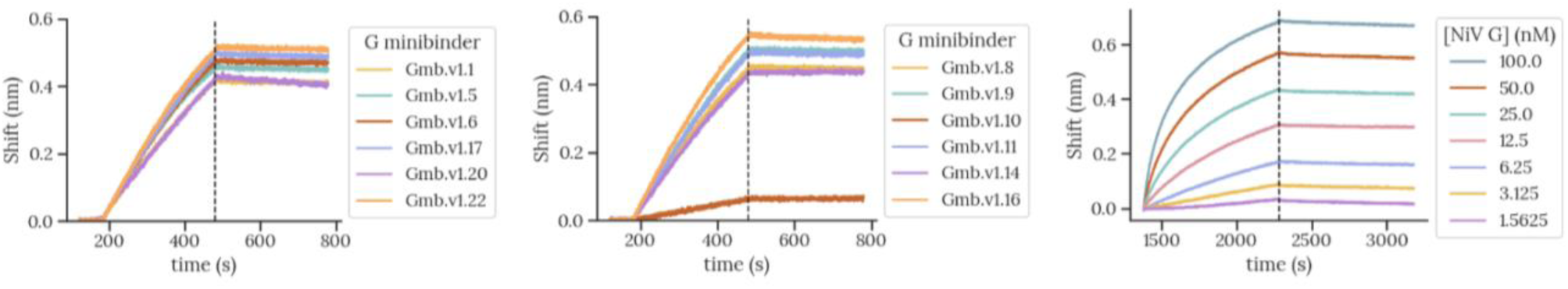
Competition BLI **(left and middle)** and multicycle kinetics (**right**) of computationally designed G minibinder v1 designs. **Competition BLI** – Fc-tagged hEphrin-B2 was immobilized onto Protein A biosensors, baseline normalized in HBS-EP(+), and then dipped into 10 nM NiV G tetramer that was pre-incubated at room temperature for 10 minutes with 1 µM SEC-purified G minibinder v1 designs (100x molar excess). Shown is the association phase (left of dashed line) and dissociation phase (right of dashed line) of NiV G tetramer in the presence of G minibinder. Minibinder alone traces (i.e. without NiV G in analyte) were subtracted from minbinder + NiV G traces to correct for nonspecific adsorption to biosensors due to high G minibinder concentrations – shown is the resultant trace. G minibinder v1.10 was the only design that promoted detectable reduction of NiV G binding signal by competition BLI. **Multicycle kinetics** – G minibinder v1.10 was immobilized via amine coupling to AR2G biosensors and NiV G tetramer was titrated down from 100 nM to 1.56 nM in a multicycle kinetics experiment to evaluate binding. Shown is the association phase (left of dashed line) and dissociation phase (right of dashed line) of NiV G tetramer.

**Supplementary Figure 3.**
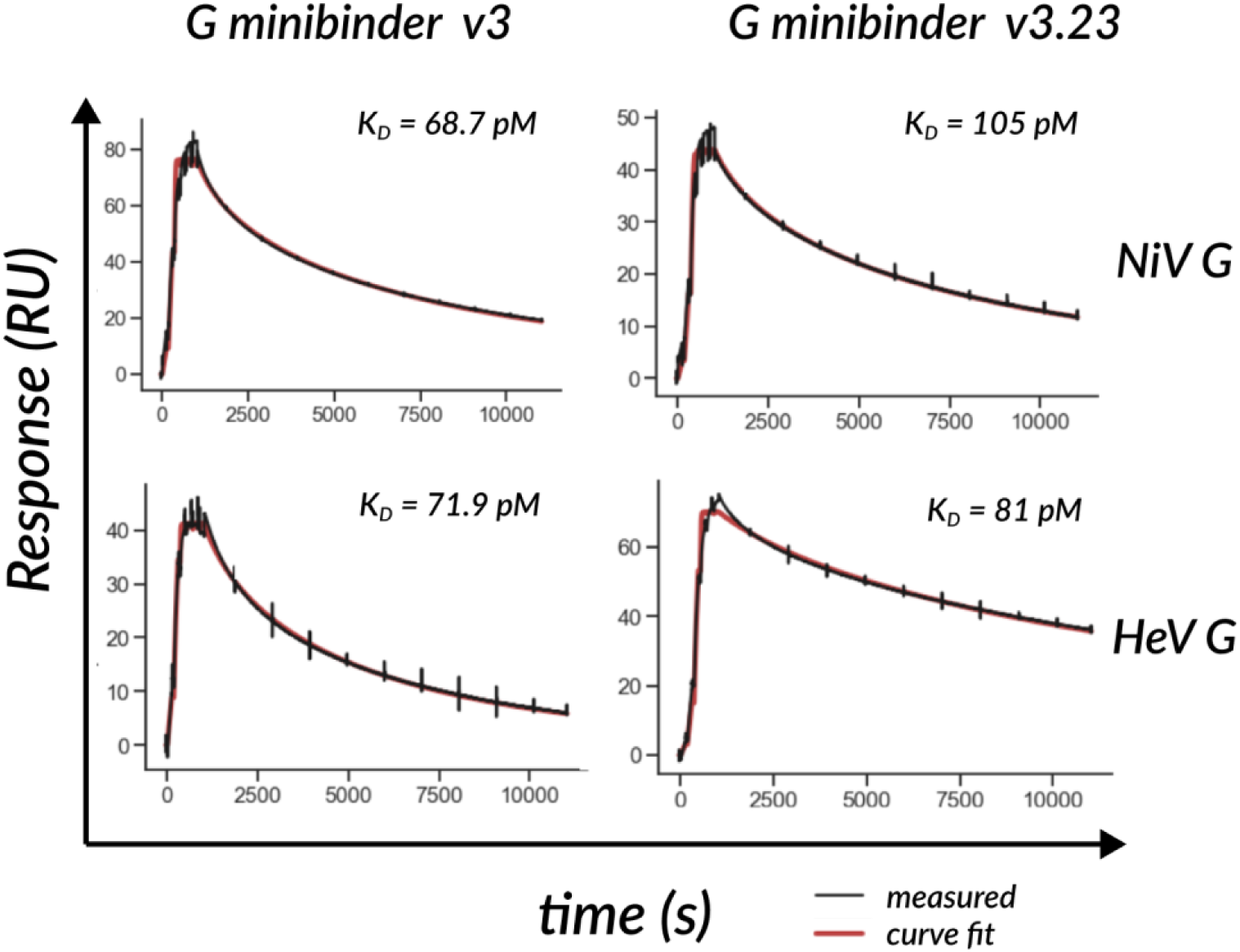
Comparative binding kinetics of anti-G v3 and v3.23 minibinders with immobilized NiV G head (amine coupling via CM5 chip). Minibinders were titrated from 5 µM to 1.6 nM over 6 x 5-fold serial dilutions. Kinetic curves (black) fit with a 1:1 binding model model (color) to extract kinetic parameters for these monovalent proteins. Though being 86% sequence similar, the anti-G minibinder v3 bound NiV G with higher affinity to both NiV and HeV G and thus was elected for further downstream development.

**Supplementary Figure 4.**
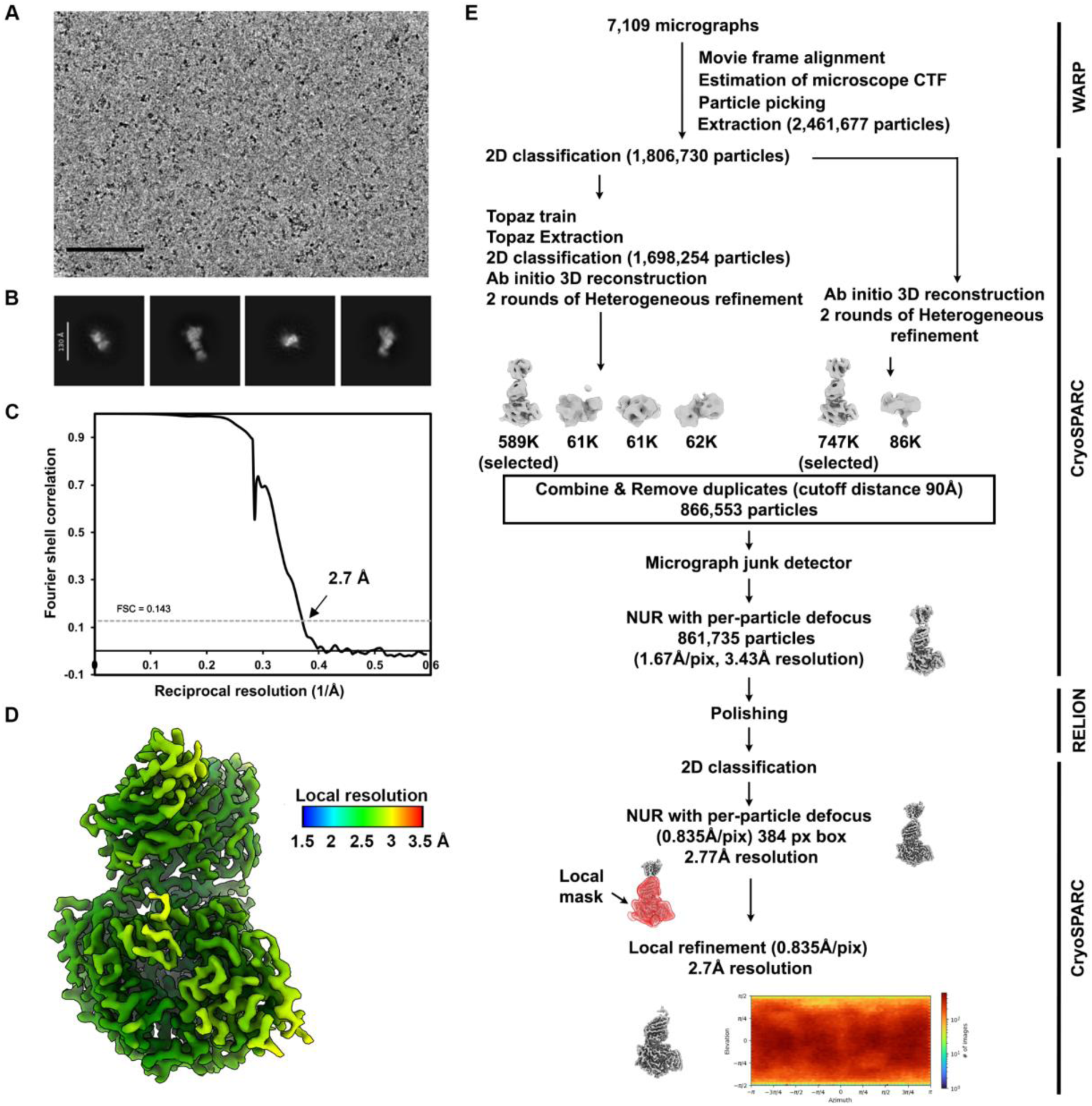
Data collection and processing of the cryo-EM dataset of the vitrified NiV G head domain bound to the G minibinder v3 and HENV32 Fab. (A-B) Representative electron micrograph and 2D class averages of the complex embedded in vitreous ice. Scale bars: 100 nm (A) and 130 Å (B). (C) Gold-standard Fourier shell correlation curve of the structure obtained by local refinement of the variable domains of the HENV32 Fab, G minibinder v3 and NiV G head. The 0.143 cutoff is indicated by a horizontal dashed line. (D) Local resolution estimation calculated using cryoSPARC and plotted on the corresponding sharpened reconstruction . (E) Data processing flowchart. CTF: contrast transfer function; NUR: non-uniform refinement. The angular distribution of particle images calculated using cryoSPARC is shown as a heat map.

**Supplementary Figure 5.**
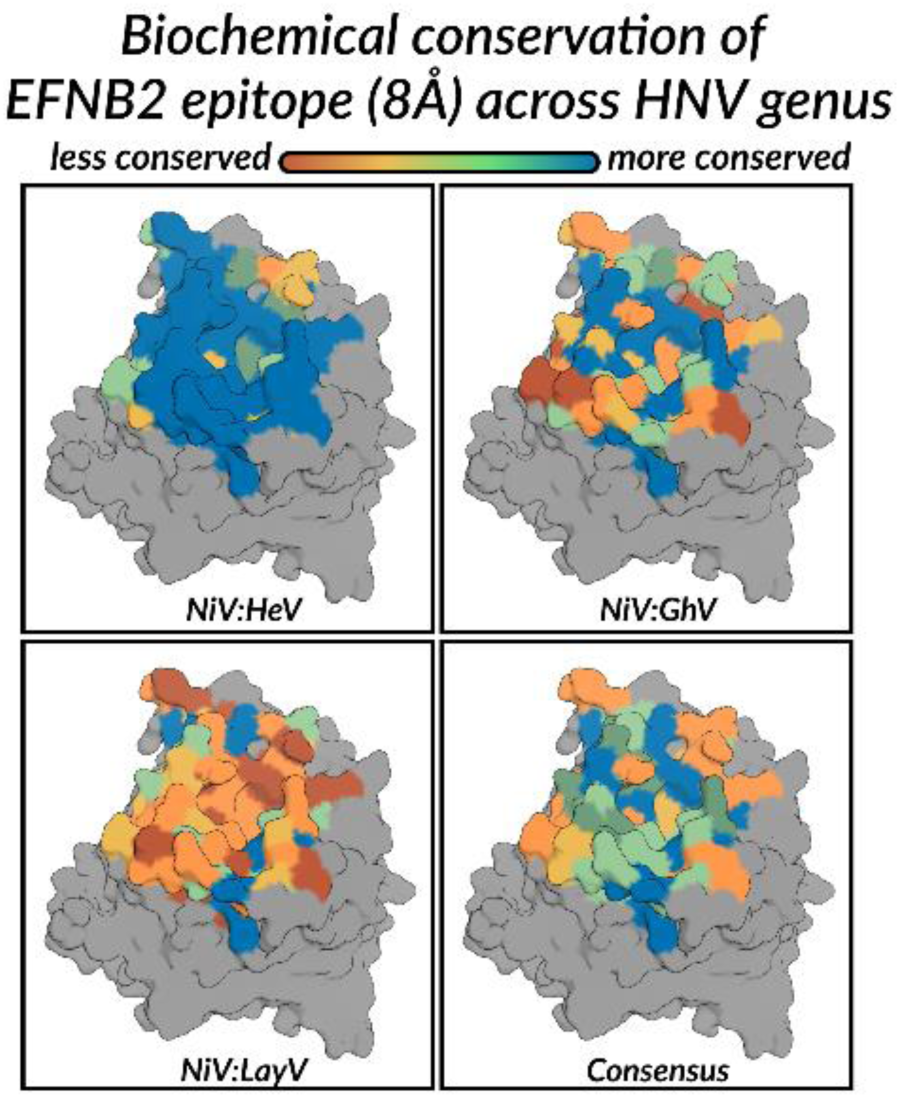
Sequence conservation of EFNB2 binding site across selected henipaviruses and parahenipaviruses colored by BLOSUM-62-weighted biochemical similarity of residues of MAFFT-aligned sequences.

**Supplementary Figure 6.**
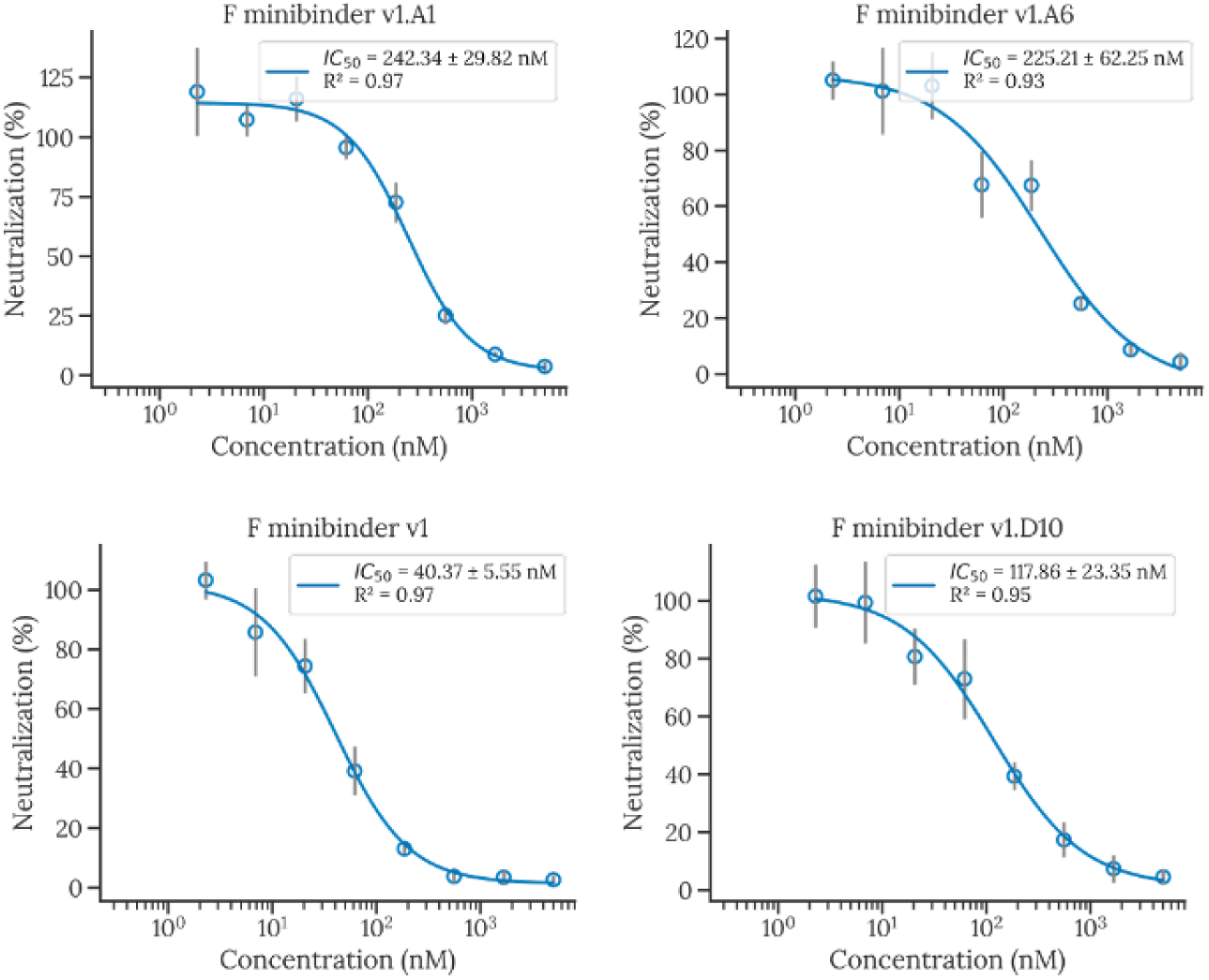
Dose-response neutralization curves of F minibinder YSD hits with detectable binding affinity to NiV F. Neutralization experiments were run with HIV pseudotypes. Biological replicates (n=4) each composed of the mean of two technical duplicates at each concentration represented as a mean (dot) and standard deviation (vertical error bar). IC_50_ and standard error of the fit reported in the legend.

**Supplementary Figure 7.**
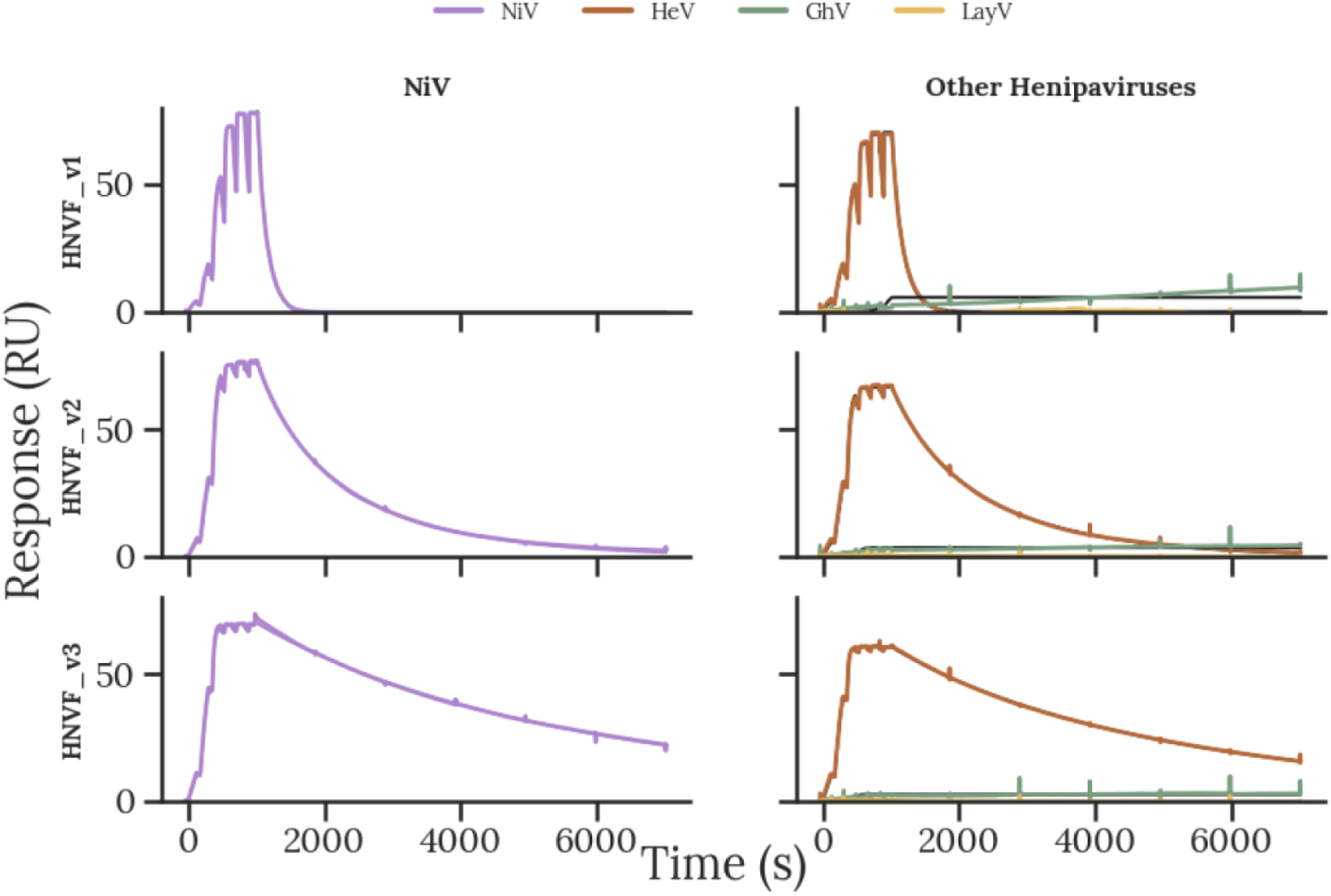
SPR sensorgrams of F minibinder v3 binding to the immobilized prefusion F ectodomain trimers (amine coupling via CM5 chip). Minibinders were titrated from 5 µM to 1.6 nM over 6 x 5-fold serial dilutions. Kinetic curves (black) fit with a 1:1 binding model model (color) to extract kinetic parameters for these monovalent proteins. Binding was observed for NiV and HeV F, but not GhV and LayV F.

**Supplementary Figure 8.**
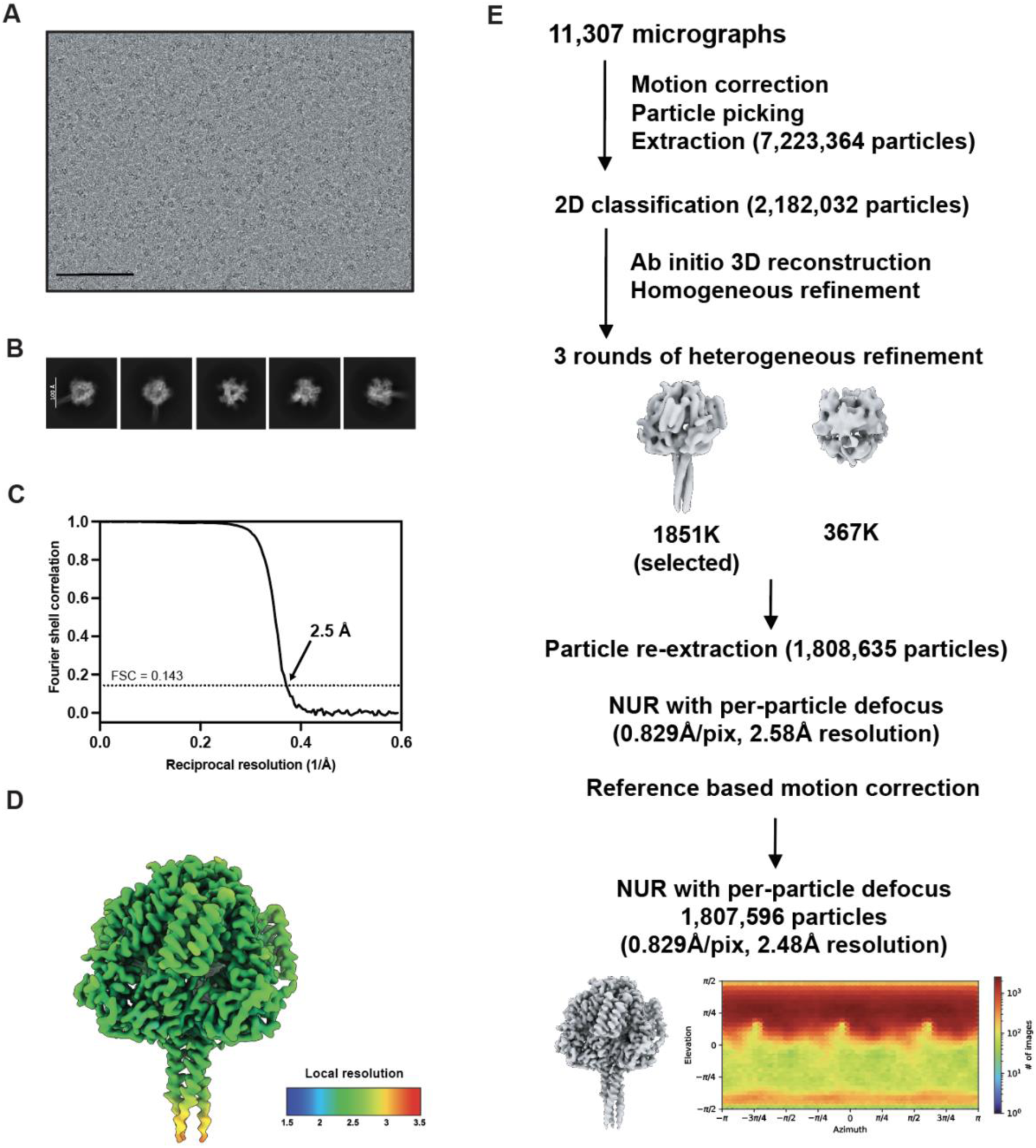
Data collection and processing of the cryo-EM dataset of the vitrified prefusion NiV F bound to the F minibinder v3. (A-B) Representative electron micrograph and 2D class averages of the complex embedded in vitreous ice. Scale bars: 100 nm (A) and 100 Å (B). (C) Gold-standard Fourier shell correlation curve of the overall reconstruction. The 0.143 cutoff is indicated by a horizontal dashed line. (D) Local resolution estimation calculated using cryoSPARC and plotted on the sharpened reconstruction. (E) Data processing flowchart. NUR: non-uniform refinement. The angular distribution of particle images calculated using cryoSPARC is shown as a heat map.

**Supplementary Figure 9.**
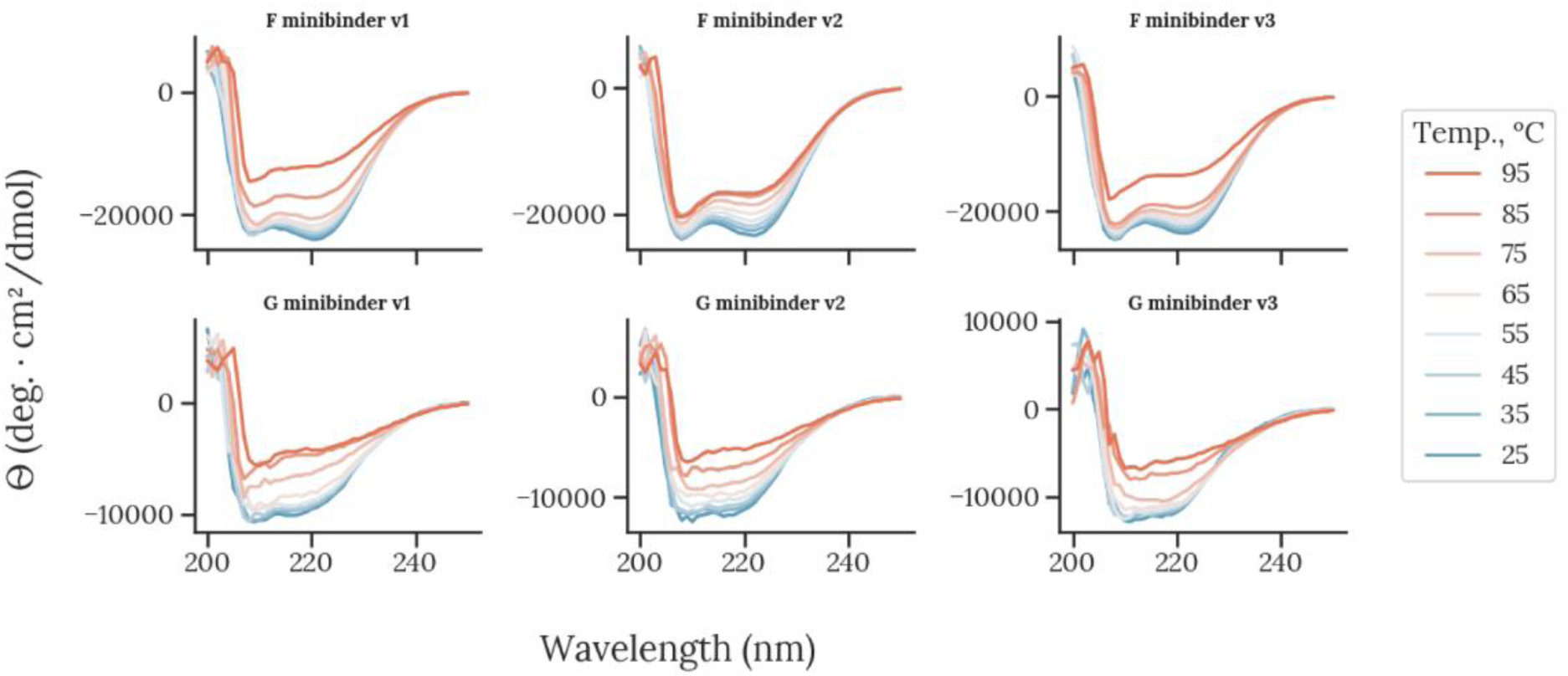
Circular dichroism spectra of F & G minibinder v1-3 monomers at temperatures ranging from 25 - 95 °C indicate only partial loss of secondary structure at 85-95 °C for all monomers.

**Supplementary Figure 10.**
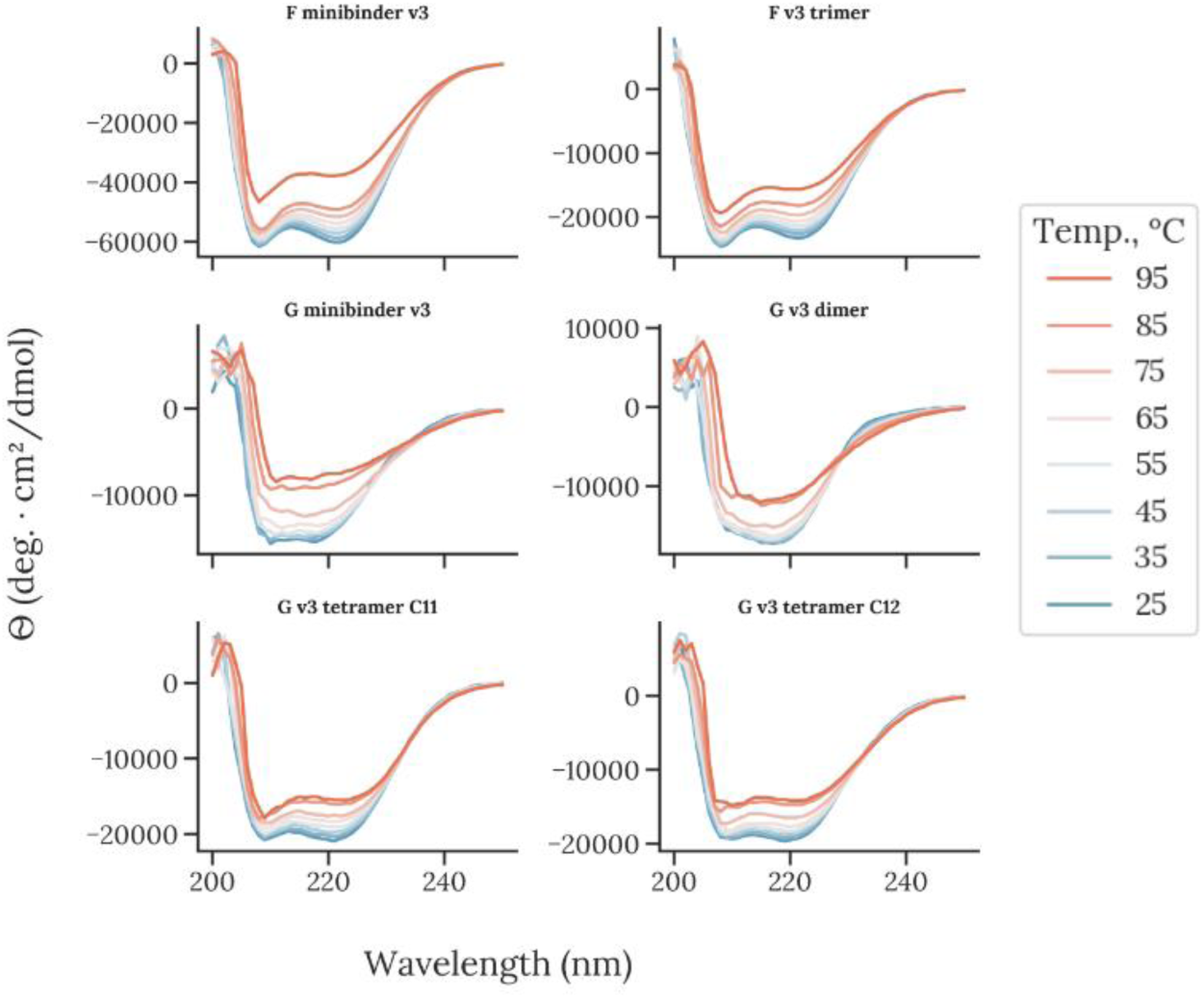
Circular dichroism spectra of F & G minibinder v3 monomers and oligomers at temperatures ranging from 25 - 95 °C indicate only partial loss of secondary structure at 85-95 °C for all constructs.

**Supplementary Figure 11.**
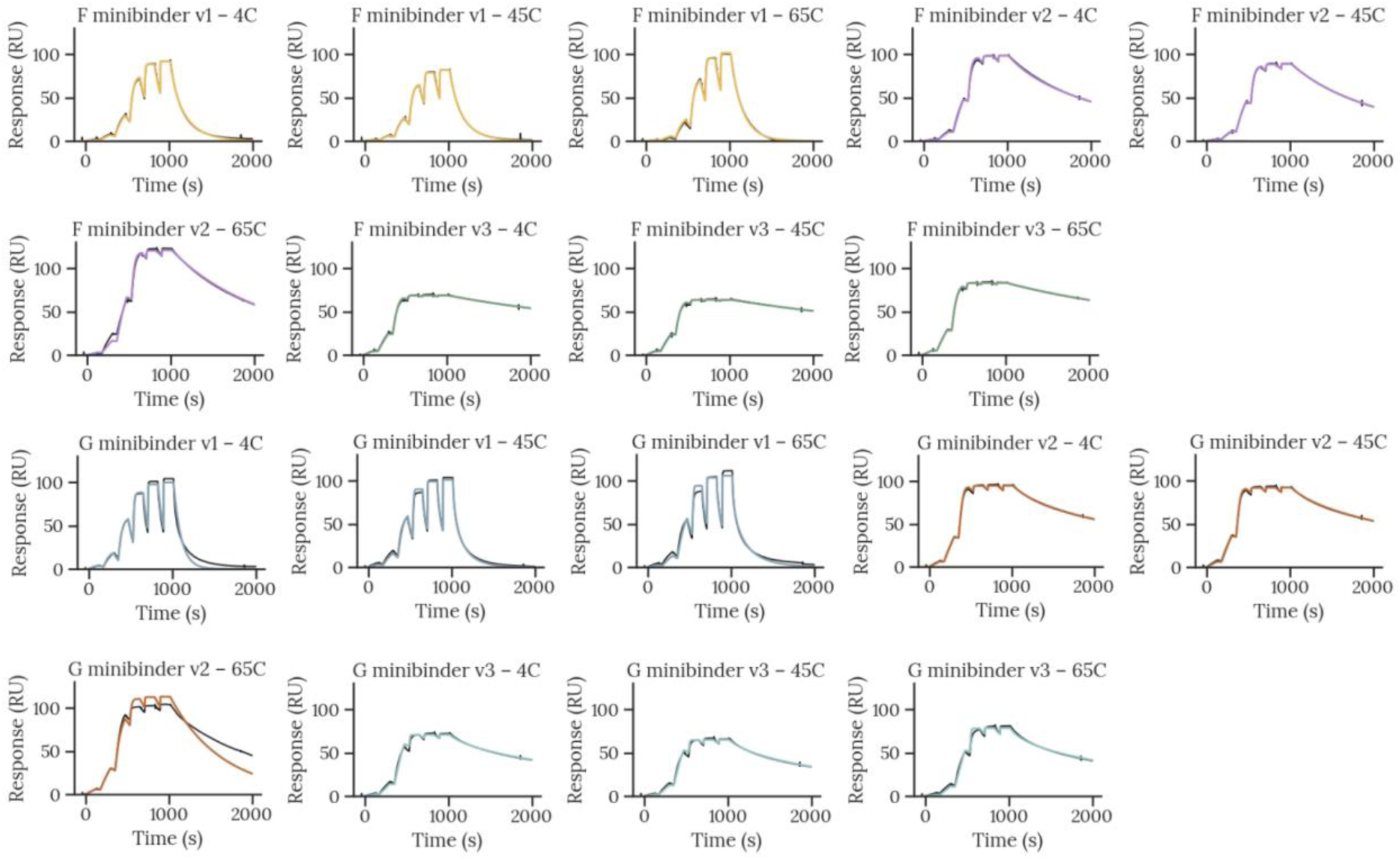
SPR sensorgrams of F & G minibinder monomers incubated for 1 week at temperatures ranging from 4 - 65°C to immobilized NiV F or G (amine coupling via CM5 chip). Minibinders were titrated from 5 µM to 1.6 nM over 6 x 5-fold serial dilutions. Kinetic curves (black) fit with a 1:1 binding model model (color) to extract kinetic parameters for these monovalent proteins. Binding is preserved for all designs across the temperature range.

**Supplementary Figure 12.**
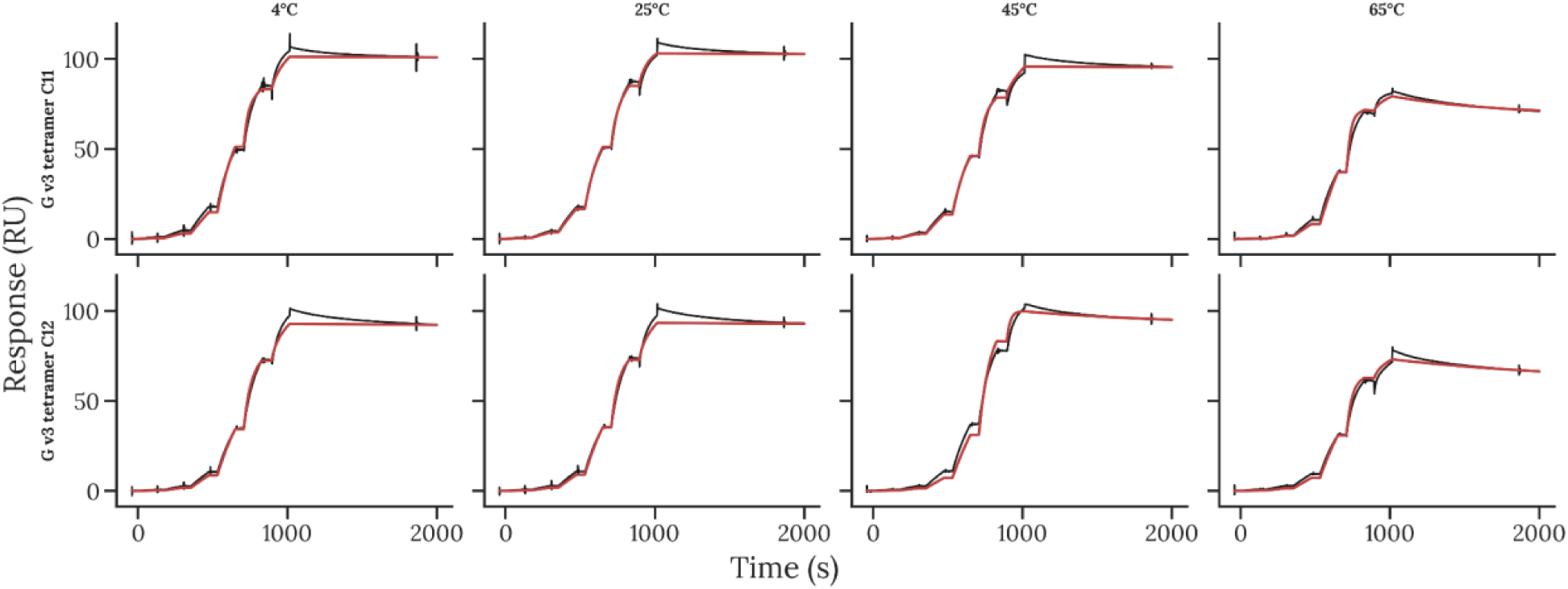
SPR kinetic curves of G minibinder v3 tetramers C11 and C12 to immobilized NiV G tetramer (amine coupling via CM5 chip) after 1-week incubation in 1x PBS at 4, 25, 45, and 65°C. Loss of peak binding signal only observed at 65°C, suggesting loss from solution via aggregation or degradation. Minibinders were titrated from 5 µM to 1.6 nM over 6 x 5-fold serial dilutions. Kinetic curves (black) fit with a bivalent analyte model (red) to account for avidity due to multivalent analyte.

**Supplementary Figure 13.**
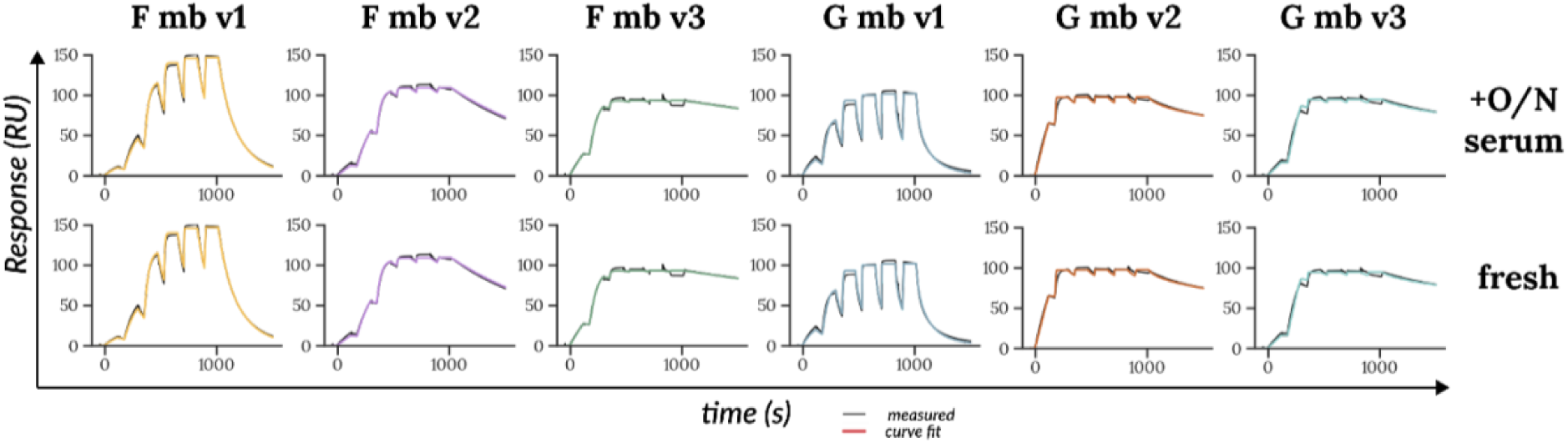
SPR binding kinetics of F & G minibinder v1-3 monomers to immobilized prefusion NiV F ectodomain trimer (amine coupling via CM5 chip) after overnight incubation of minibinders in human serum at 37°C vs fresh addition to human serum prior to run. Minibinders were titrated from 5 µM to 1.6 nM over 6 x 5-fold serial dilutions. Kinetic curves (black) fit with a 1:1 binding model model (color) to extract kinetic parameters for these monovalent proteins.

**Supplementary Figure 14.**
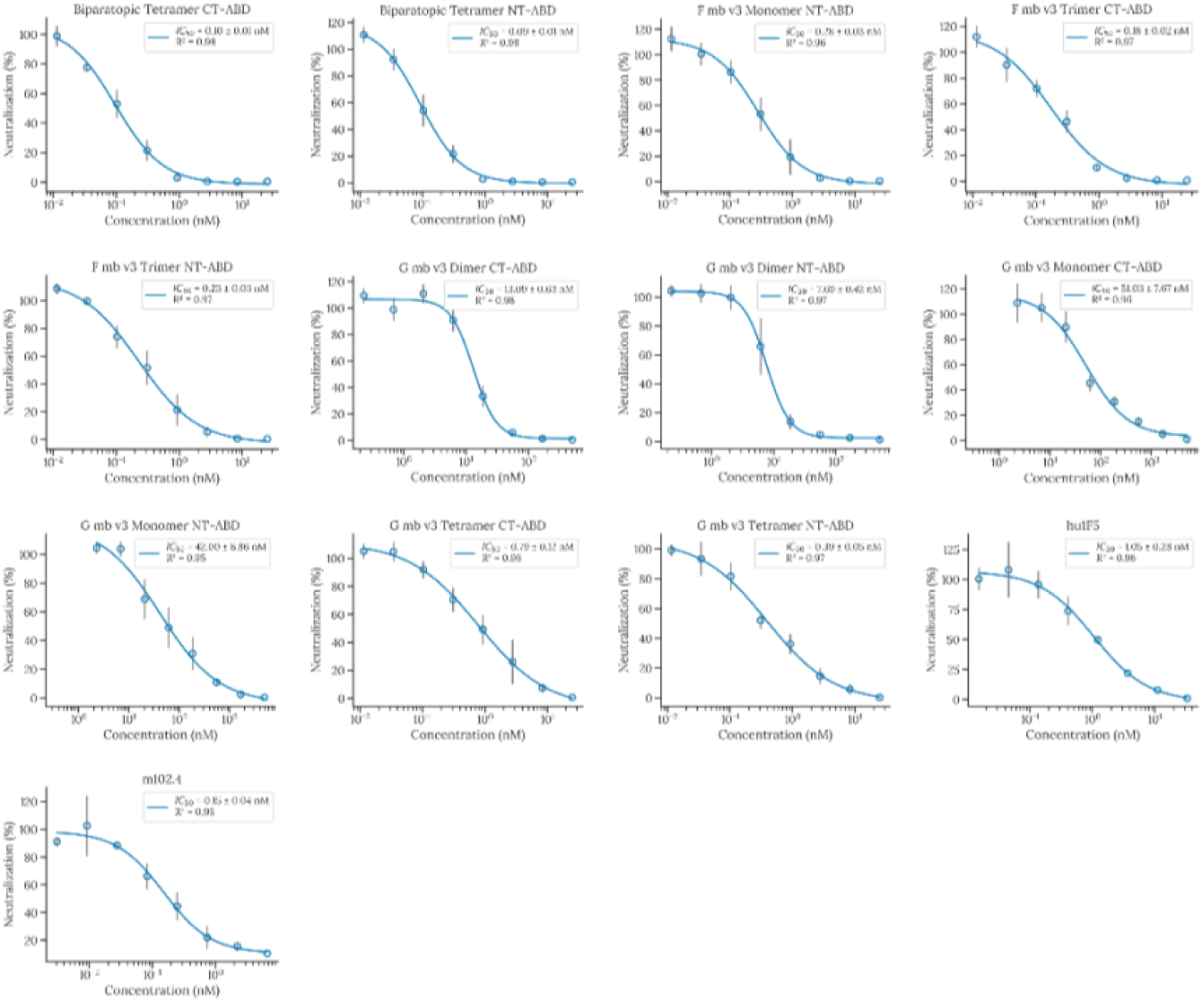
Dose-response neutralization curves of F + G minibinder v3 monomers and oligomers compared with mAbs hu1F5 (anti-HNV F) and m102.4 (anti-HNV G). Technical replicates (n=4) at each concentration represented as a mean (dot) and standard deviation (vertical error bar). IC_50_ and standard error of the fit reported in the legend.

**Supplementary Figure 15.**
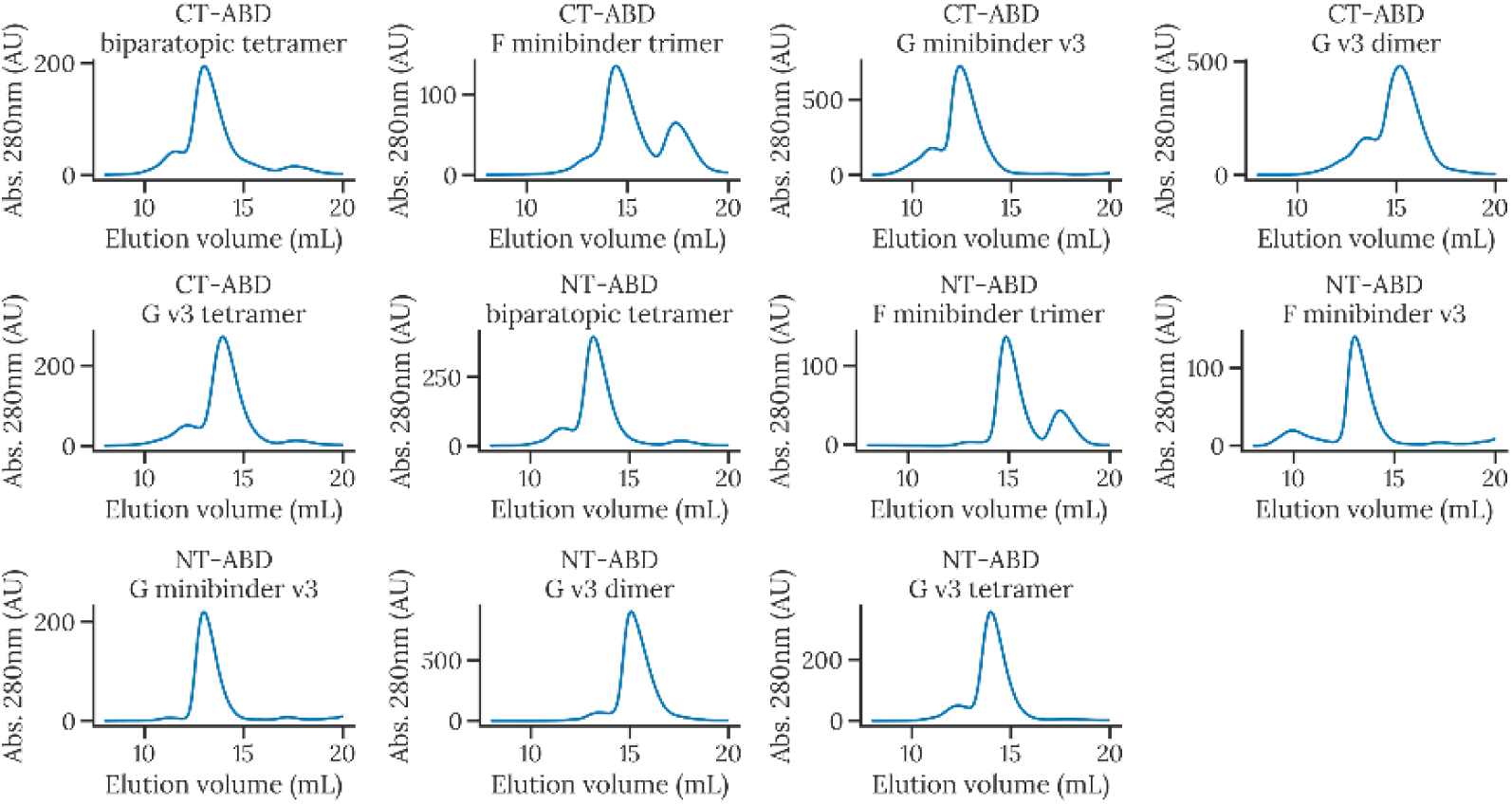
Size exclusion chromatograms using Superdex S75 (for monomeric minibinders) or S200 (for oligomers) 10-300 GL Increase for v3 monomers and oligomers conjugated to albumin-binding domains recombinantly purified from *E. coli* and eluted in phosphate-buffered saline, pH 7.4.

**Supplementary Table 1.** SPR kinetics and NiV F & G-pseudotyped HIV IC_50_ values of F minibinder v1 designs.

| Design name | NiV F K<sub>D</sub> (M) | NiV F k<sub>a</sub> (1/Ms) | NiV F k<sub>d</sub> (1/s) | HeV F K<sub>D</sub> (M) | HeV F k<sub>a</sub> (1/Ms) | HeV F k<sub>d</sub> (1/s) | IC<sub>50</sub> (nM) | IC<sub>50</sub> std. dev. (nM) | R<sup>2</sup> |
| --- | --- | --- | --- | --- | --- | --- | --- | --- | --- |
| F mb v1.A1 | 1.21E-07 | 217000 | 0.0262 | 1.04E-07 | 260000 | 0.0272 | 242.3373 | 29.8159 | 0.966217 |
| F mb v1.A6 | 1.19E-07 | 599000 | 0.0712 | 7.68E-08 | 618000 | 0.0474 | 225.2061 | 62.24712 | 0.931804 |
| F mb | 1.55E- | 462000 | 0.007 | 4.34E | 45800 | 0.019 | 117.8 | 23.35119 | 0.949 |
| v1.D10 | 08 |  | 16 | -09 | 00 | 9 | 649 |  | 69 |
| F mb<br>v1.A7<br>(v1) | 7.43E<br>-08 | 604000 | 0.044<br>9 | 3.5E-08 | 73800<br>0 | 0.025<br>8 | 40.37<br>356 | 5.546531 | 0.968<br>555 |

**Supplementary Table 2.** IC_50_ values of ABD-conjugated F and G minibinder v3 monomers and oligomers tested in NiV F & G-pseudotyped HIV assay.

| **Design name** | **IC<sub>50</sub>** | **IC<sub>50</sub> std. dev.** | **R<sup>2</sup>** |
| --- | --- | --- | --- |
| G mb v3 Monomer NT-ABD | 41.99717 | 8.863075 | 0.951728 |
| F mb v3 Trimer CT-ABD | 0.175589 | 0.021899 | 0.968035 |
| G mb v3 Monomer CT-ABD | 51.03011 | 7.674627 | 0.955398 |
| Biparatopic Tetramer CT-ABD | 0.096955 | 0.008909 | 0.981219 |
| G mb v3 Dimer CT-ABD | 13.08776 | 0.633395 | 0.980098 |
| F mb v3 Monomer NT-ABD | 0.279482 | 0.02757 | 0.964669 |
| G mb v3 Tetramer NT-ABD | 0.389201 | 0.050227 | 0.969691 |
| G mb v3 Tetramer CT-ABD | 0.786113 | 0.121969 | 0.960881 |
| G mb v3 Dimer NT-ABD | 7.686403 | 0.424577 | 0.971988 |
| Biparatopic Tetramer NT-ABD | 0.09189 | 0.007162 | 0.981786 |
| F mb v3 Trimer NT-ABD | 0.228445 | 0.025805 | 0.97351 |
| m102.4 | 0.152094 | 0.040877 | 0.950154 |

